# PKA can adopt structural or enzymatic roles depending on its anchoring site

**DOI:** 10.64898/2026.08.12.743961

**Authors:** Timothy W. Church, Yuelin Luo, Ryan S. Dowsell, Ian J. White, Matthew G. Gold

## Abstract

Protein kinase A (PKA) is a central mediator of cAMP signaling whose spatial specificity is conferred by A-kinase anchoring proteins (AKAPs). However, functions of individual anchored PKA pools are difficult to resolve as current approaches cannot isolate anchoring at defined sites. Here, we develop a SpyCatcher-SpyTag-based replacement strategy enabling direct comparison of AKAP function with and without type II PKA regulatory (RII) subunit anchoring in intact signaling environments. Anchoring RII to AKAP79 or AKAP1 produced distinct phosphorylation-dependent effects, including catalytic subunit membrane recruitment and mitochondrial morphology regulation. In contrast, MAP2-anchored RII drove phosphorylation-independent microtubule straightening, increased formation of widely spaced bundles, and supported dendritic arborization. This structural function was recapitulated by native MAP2-RII binding and required an intact cAMP-binding pocket in the second cyclic nucleotide-binding domain. These findings establish that AKAP-bound PKA can function as either an enzymatic or structural component and provide a generalizable strategy to dissect compartmentalized kinase signaling.

## Introduction

cAMP-dependent protein kinase, also known as protein kinase A (PKA), is the major intracellular receptor for cAMP^1^. PKA phosphorylation underlies numerous biological responses including sympathetic stimulation of the heart^2^, synaptic strengthening^3^, and glycogen breakdown^4^. A long-standing question in cell signaling is how PKA is directed to phosphorylate specific subsets of protein substrates in a cell-type- and stimulus-dependent manner since the kinase exhibits relatively broad substrate selectivity in the test tube. A model has emerged with A-kinase Anchoring Proteins (AKAPs) playing a key role by positioning PKA at different sub-cellular locations and limiting PKA phosphorylation to substrates in their locale^5^. AKAPs present ∼20 amino acid amphipathic helices that dock to the N-terminal dimerization and docking (D/D) domain of PKA regulatory subunits^6,7^ with a general preference for type II (RII) rather than type I (RI) regulatory subunits. Each AKAP also contains a targeting sequence. For example, polybasic regions and palmitoyl groups target AKAP79 to the cell membrane^8^, MAP2 binds microtubules via C-terminal tubulin-binding motifs^9^, and AKAP1 contains an N-terminal sequence that inserts into the outer mitochondrial membrane^10^. The canonical AKAP model posits that tetrameric PKA holoenzymes comprised of two regulatory and two catalytic (C) subunits dock to each AKAP site, with local rises in cAMP triggering C subunit release. However, recent discoveries show both that PKA C subunits are greatly outnumbered by RII subunits^11^, and that dephosphorylation of RII subunits is also an important factor that dictates how quickly free C subunits are captured^12^. In this study, we aimed to make sense of these new aspects of cAMP signaling by isolating the effects of individual PKA anchoring sites using a molecular glue technology.

Quantitative immunoblotting^13,14^ and proteomics^15^ have shown PKA regulatory subunits to be in large excess of C subunits in all tissues tested. In HEK293T cells – the leading model for studying cAMP signaling – both RI and RII subunits substantially exceed C subunits (∼1.5 μM RII, 0.6 μM RI, 0.2 μM C). In brain tissues, combined PKA regulatory subunits typically exceed C subunits by more than 20-fold^13^. Such a pronounced excess suggests that PKA regulatory subunits may serve roles beyond simple sequestration of C subunits. While such functions have not yet been defined, recent work on other highly abundant neuronal signaling proteins, including SynGAP and CaMKII, has shown that enzymatic activity can coexist with — or be secondary to — broader, non-enzymatic roles^16,17^. A further complication arises from the phosphorylation-dependent regulation of RII subunits, which modulates their capacity to capture C subunits. Unlike RI, RII subunits are phosphorylated within the inhibitory sequence by bound C subunits, and this phosphorylated (‘pRII’) state often persists following C subunit release^12^. Surface plasmon resonance measurements indicate that dephosphorylated RII captures C subunits approximately 50-fold faster than pRII^12^, yet free pRII is a poor substrate for most cellular phosphatases. Importantly, AKAP79 has been shown to facilitate efficient dephosphorylation of pRII by calcineurin (CN), which it co-anchors alongside RII^14^. Given the large excess of RII over C subunits, these findings raise the possibility that C subunit occupancy is not uniform, but instead varies between anchoring sites, with local phosphatase activity influencing the rate and extent of C subunit capture. Testing these ideas, however, remains challenging with existing experimental approaches.

Studies of nanodomain cAMP signaling have benefited from availability of peptide disruptors that target R subunit docking sites^18^, with synthetic disruptors now available for RI and RII selective disruption^6,19^ or with internal staples for enhanced stability^20^. However, it has proved to be challenging to develop AKAP-selective binding sequences, which would help to decipher the functions of individual binding sites^21^. In this study, we took a different tack by using the SpyCatcher-SpyTag molecular glue technology to dictate where RII is anchored in cells. The SpyCatcher-SpyTag system is derived from the *Streptococcus pyogenes* fibronectin-binding protein^22^, and comprises a SpyCatcher domain that forms a highly-stable isopeptide bond with the central aspartate of the 13-amino acid SpyTag sequence^23^. The SpyTag sequence is a similar size to AKAP anchoring helices (**Fig. 1A**), and can be introduced at internal sites without affecting conjugation efficacy^24^. After confirming that this system can efficiently substitute for native RII– AKAP anchoring, we developed cell lines that enable delineation of AKAP-selective type II PKA anchoring functions. By enabling selective control of RII anchoring, this approach allows us to examine how local anchoring environments – including phosphatase access – shape C subunit occupancy and support functions beyond local phosphorylation.

**Figure 1.**
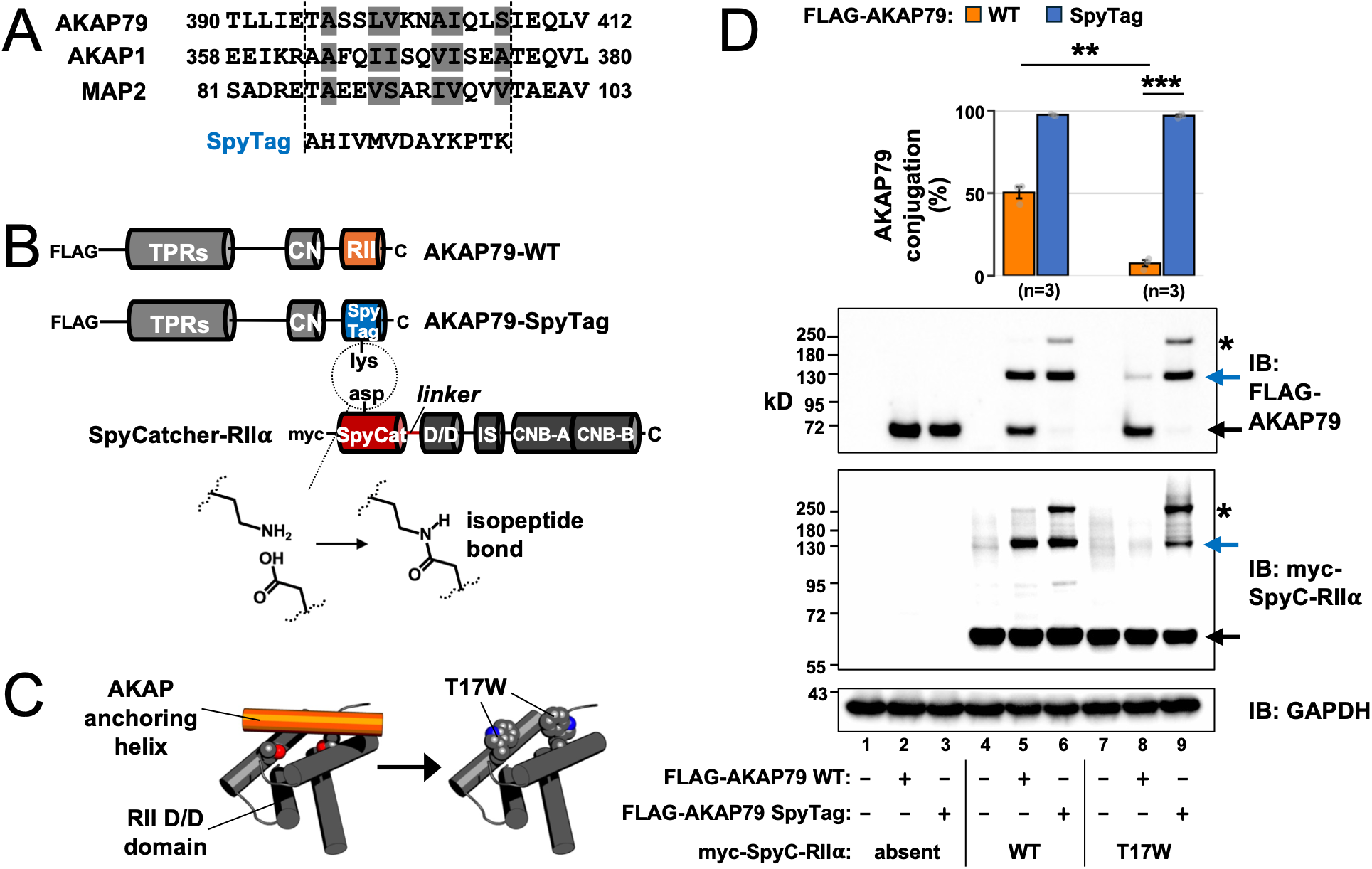
Isopeptide-based RIIα anchoring to AKAP79 via the SpyCatcher-SpyTag system. (**A**) Alignment of RII anchoring helices (core aliphatics in grey). Dotted lines indicate location of SpyTag substitutions. (**B**) AKAP79 and RII expression constructs, showing locations of SpyTag sequence (blue) in place of the native anchoring helix (orange) in AKAP79, and SpyCatcher domain (red) at the RIIα N-terminus for isopeptide linking with AKAP79-SpyTag. The inhibitor sequence (IS), and tandem cyclic nucleotide binding (CNB) domains of RII are shown. (**C**) Location of the T17W mutation within the RIIα D/D domain (grey) that sterically blocks binding to AKAP anchoring helices (orange). The basic structure is PDB ID 2IZX. (**D**) Immunoblots of HEK293T lysates following transfection with different combinations of RIIα and AKAP79 constructs, as shown. Unconjugated proteins (black arrows), 1:1 AKAP79:RIIα conjugates (blue arrows), and likely 1:2 AKAP79:RIIα conjugates (asterisks) are indicated. Conjugation efficiency (top sub-panel) was calculated by densitometry of anti-FLAG-AKAP79 IBs with data analyzed using unpaired two-tailed Student’s t tests (***p < 0.001, **p < 0.01).

## Results

### RII subunit anchoring can be mediated through SpyCatcher-SpyTag isopeptide linking

We first aimed to test whether the SpyCatcher-SpyTag system could be used as a substitute for the PKA-AKAP interface. We developed a vector for mammalian expression of RIIα with the minimal SpyCatcher domain^25^ fused at its N-terminus (red, **Fig. 1B**), alongside vectors for expression of either wild-type (WT) FLAG-tagged AKAP79, or a variant in which the central 13 amino acids of its anchoring helix were replaced with the SpyTag sequence – ‘AKAP79-SpyTag’ (blue, **Fig. 1B**). We also generated a variant of SpyCatcher-RIIα that includes the T17W substitution, which disrupts docking to native anchoring helices (**Fig. 1C**)^21^. We compared conjugation following co-transfection of different combinations in HEK293T cells (**Fig. 1D**).

No band at ∼140 kD, corresponding to the expected AKAP79-RII conjugate, was detected when either AKAP79 variants (lanes 2–3, **Fig. 1D**) or SpyCatcher-RIIα variants (lanes 4 and 7) were expressed alone. Co-expression of myc-SpyCatcher-RIIα variants with either the wild-type (WT) (lane 5) or the SpyTag variant of AKAP79 (lane 6) generated a prominent band at ∼140 kD indicative of 1:1 AKAP79-RII conjugation. Comparison of anti-FLAG (top panel, **Fig. 1D**) and anti-myc (middle panel) immunoblots (IBs) shows that SpyC-RIIα is in large excess of AKAP79 in all co-expression conditions. Therefore, to compare the efficiency of conjugation between conditions, we used densitometry to quantify the relative levels of AKAP79 in 1:1 AKAP79-RII conjugates (indicated by blue arrow in top panel of **Fig. 1D**) compared to unconjugated AKAP79 (black arrow). Surprisingly, we found that approximately half of AKAP79-WT formed conjugates with SpyCatcher-RIIα (50.3 ± 3.6 %, orange bars, **Fig. 1D**). When the SpyTag was substituted into AKAP79, conjugation efficiency approached completion (97.3 ± 0.3 %, blue bars, **Fig. 1D**). This shows that conjugation to SpyTag-AKAP79 is highly efficient but that non-specific isopeptide bond formation also occurs regardless of whether the SpyTag sequence is present. The native anchoring helix in AKAP79 binds to the D/D domain of RIIα with sub-nanomolar affinity^26^. We hypothesized that this interface was enabling SpyCatcher-RIIα to form isopeptide bonds to AKAP79-WT by increasing the effective concentration of aspartate residues present close to the anchoring helix (there are 4 aspartates within 20 amino acids of the RII anchoring helix). Consistent with this notion, incorporation of the T17W substitution into SpyCatcher-RIIα reduced isopeptide bond formation to FLAG-AKAP79-WT (lane 8, **Fig. 1D**) to 7.4 ± 2.0 % (*p* = 0.0016) but did not alter efficient conjugation to AKAP79-SpyTag (96.8 ± 0.8 %, lane 9, **Fig. 1D**). A higher molecular weight (MW) band at ∼ 250 kD (asterisks, **Fig. 1D**) immunoreactive for both RII and AKAP79 was also prominent in cells co-expressing SpyCatcher-RIIα variants and AKAP79-SpyTag, which is likely to correspond to a covalently-linked trimer with two copies of RII and one copy of AKAP79. In sum, our data show that fusing the SpyCatcher domain to the N-terminus of RIIα T17W enables efficient and specific isopeptide linking to AKAP79 bearing the SpyTag sequence in place of its RII anchoring helix.

### Development of a SpyCatcher-RII cell line that facilitates AKAP-selective PKA anchoring

We next set out to develop a cell line that would enable us to exploit SpyCatcher-SpyTag linking for selective PKA anchoring. We focused on HEK293T cells, which continue to be the leading model cell line for understanding nanodomain cAMP signaling^27,28^. In WT HEK293T cells, PKA is anchored at multiple sites (**Fig. 2A**) including AKAP79, MAP2, and AKAP1^29^. We aimed to develop a cell line that expresses only SpyCatcher-RIIα T17W subunits, which are not anchored (**Fig. 2B**) unless a specific SpyTag-substituted AKAP is introduced (**Fig. 2C**). We first knocked out both endogenous RII isoforms using CRISPR-Cas9 targeting to generate ‘ΔRII’ cells before introducing myc-SpyCatcher-RIIα T17W subunits into these ΔRII cells by lentiviral integration (**Fig. 2D**). Immunoblotting confirmed effective knockout of RIIα and RIIβ in ΔRII cells (**Fig. 2E**) and identified a clone expressing myc-SpyCatcher-RIIα T17W at comparable levels to endogenous RIIα in WT HEK293T cells (clone 1, **Fig. 2E**). This clone, hereafter referred to as ‘SpyC-RII’, was taken forward for all subsequent experiments. Additional immunoblots revealed no obvious compensatory changes in RI or C subunit expression in either ΔRII or SpyC-RII cell lines (**Fig. S1**).

**Figure 2.**
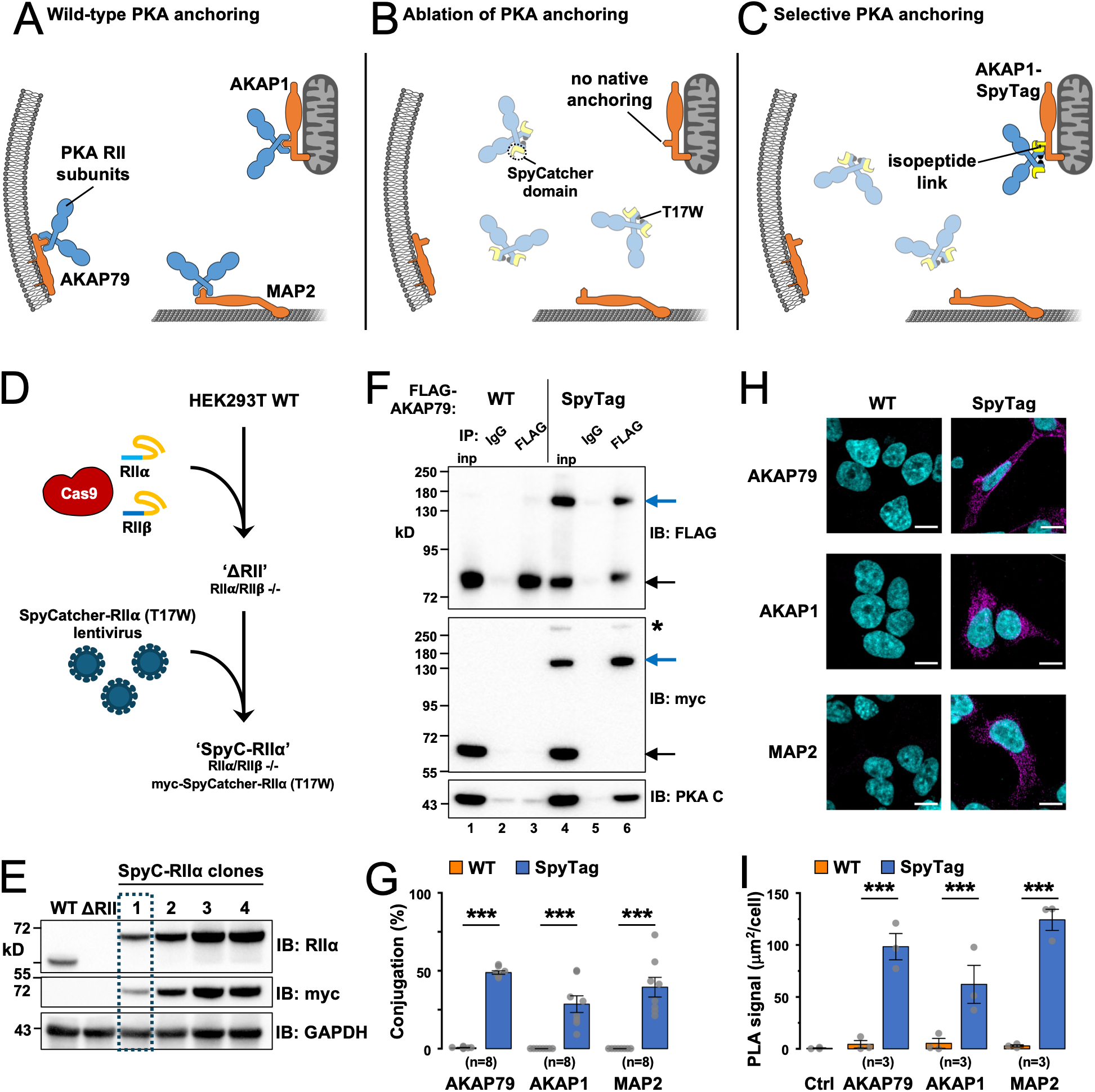
Development of a cell line for AKAP-selective isopeptide PKA anchoring. (**A**) Representation of RII subunit anchoring at three example sites in WT HEK293T cells. In cells where RII subunits are replaced with SpyCatcher-RIIα (T17W) subunits, anchoring to endogenous AKAPs does not occur (**B**), but selective anchoring (**C**) can be achieved through expression of a SpyTag-substituted AKAP of choice, e.g., AKAP1 as shown. (**D**) Workflow for developing ΔRII and SpyC-RII variant HEK293T cells. (**E**) Immunoblots showing expression of RIIα and myc-SpyCatcher-RIIα (T17W) subunits in clonal cell lines, including four possible SpyC-RII cell clones (clone 1 was taken forward). (**F**) Anti-FLAG or control IgG IPs from SpyC-RII cells expressing either WT or SpyTag variants of FLAG-AKAP79. Co-IP of myc-SpyCatcher-RIIα (T17W) and endogenous PKA C subunits was detected by IB. Unconjugated (black) and conjugated (blue) proteins are indicated with arrows. (**G**) Conjugation efficiencies for three AKAPs according to densitometry of anti-FLAG IBs of the type shown in the preceding panel. (**H**) Representative PLA images of SpyC-RII cells transfected with either WT (left column) or SpyTag (right column) variants of three AKAPs. The scale bar corresponds to 20 µm. (**I**) Quantification of PLA signal in the six conditions shown in the preceding panel. For panels G and I, statistical comparisons were performed using unpaired two-tailed Student’s t-tests (\*\*\**P* < 0.001).

We aimed to use the SpyC-RII cells to dissect the functions of specific RII anchoring sites by comparing cell phenotypes following transfection with either the WT or SpyTag-substituted variants of different AKAPs. Differences between the WT and SpyTag variants would reveal functions specific to only that RII anchoring site. Before applying this approach, we first sought to confirm that the system was working as intended. We focused on AKAP79, AKAP1 and MAP2. Transfection of SpyC-RII cells with SpyTag-AKAP79 led to formation of a band at ∼140 kD corresponding to conjugation of the AKAP and SpyCatcher-RII subunits (lane 4, **Fig. 2F**) whereas there was no detectable AKAP-RII conjugate in cells expressing WT AKAP79. This is an improvement on pilot experiments with highly overexpressed SpyCatcher-RII subunits (**Fig. 1D**). PKA C subunits were detected in anti-FLAG immunoprecipitates from cells expressing SpyTag but not WT AKAP79 (lanes 3 & 6, **Fig. 2F**), showing that the presence of the SpyTag leads to recruitment of both RII and C subunits to AKAP79. Similar results were obtained for AKAP1 and MAP2 (**Fig. S2**). Conjugation was readily detectable for all three SpyTag-substituted AKAPs, with efficiencies ranging from 28.6 ± 5.3% to 48.9 ± 1.1%, whereas all WT variants remained at baseline levels (**Fig. 2G**).

As an *in situ* test of AKAP-RII conjugation efficiency and selectivity, we turned to proximity ligation assays (PLA). We compared PLA puncta formation – indicative of proteins within 40 nm nm^30^ – in SpyC-RII cells expressing either WT or SpyTag variants of AKAP79, AKAP1, and MAP2. PLA was performed by pairing anti-myc (SpyCatcher-RII) and anti-FLAG (AKAP) antibodies. EGFP was co-transfected with FLAG-AKAPs to enable calculation of average PLA signal area per transfected cell with mean values calculated from three independent batches of cells. PLA signal was much higher in cells expressing SpyTag-AKAPs, as expected (**Fig. 2H**). PLA signal was at low levels for cells expressing either empty vector (0.36 ± 0.13 µm^2^ per cell), or WT variants of AKAP79 (3.4 ± 1.4 µm^2^), AKAP1 (5.7 ± 1.4 µm^2^), or MAP2 (3.2 ± 1.28 µm^2^). PLA puncta were markedly increased for all SpyTag variants, rising to 96.5 ± 10.7 µm^2^ per cell for AKAP79, 58.4 ± 5.9 µm^2^ for AKAP1, and 125.5 ± 16.5 µm^2^ for MAP2 (**Fig. 2I**, *P* < 0.001 for all three SpyTag versus WT comparisons). In sum, our data show that the SpyC-RII cell line provides an efficient platform for AKAP-selective anchoring of PKA RII subunits.

### RII anchored to AKAP79 acts in concert with CN to concentrate catalytic subunits at the cell membrane

We aimed to take advantage of the SpyC-RII cell line to investigate PKA anchoring at three prototypic AKAPs, beginning with AKAP79. In addition to membrane-tethering elements, AKAP79 contains a ‘PIAIIIT’ CN anchoring sequence within 50 amino acids of its PKA anchoring helix. CN can efficiently dephosphorylate pRII subunits but only when the two proteins are co-anchored to AKAP79^14^. We have previously shown *in vitro* that co-anchoring of CN and RII by AKAP79 supports faster PKA C subunit recapture via dephosphorylated RII subunits within the AKAP79 complex^14^. Before using SpyC-RII cells to explore these ideas in a more physiological context, we first tested whether anchoring to AKAP79 localized PKA C subunits as expected. We employed a vector expressing PKA Cβ bearing EGFP at its C-terminus via a 14-amino acid polyglycine linker (‘C-GFP’). Similar constructs have been used previously for live imaging of C subunit dynamics in neurons^31,32^. In all cases, the C-GFP vector was included at 1 part in 100 of DNA in transfection mixes, to limit changes to the R:C subunit ratio. We compared the distribution of C-GFP subunits in SpyC-RII cells transfected with either WT (**Fig. 3A**) or SpyTag (**Fig. 3B**) AKAP79, imaging live with the cell membrane stain wheat-germ agglutinin CF633 (WGA-CF633). Upon co-expression with WT AKAP79, which cannot anchor SpyC-RII subunits, C-GFP was distributed evenly throughout the cell as expected (**Fig. 3A**). Co-expression with AKAP79-SpyTag led to marked C-GFP enrichment at the cell membrane (**Fig. 3B**), with clear colocalisation of C-GFP (cyan, **Fig. 3B**) and WGA-CF633 (magenta) captured by a higher Pearson’s correlation between C-GFP and WGA (0.72 ± 0.02) than in cells expressing WT AKAP79 (0.19 ± 0.04, *P* < 0.001, **Fig. 3C**).

**Figure 3.**
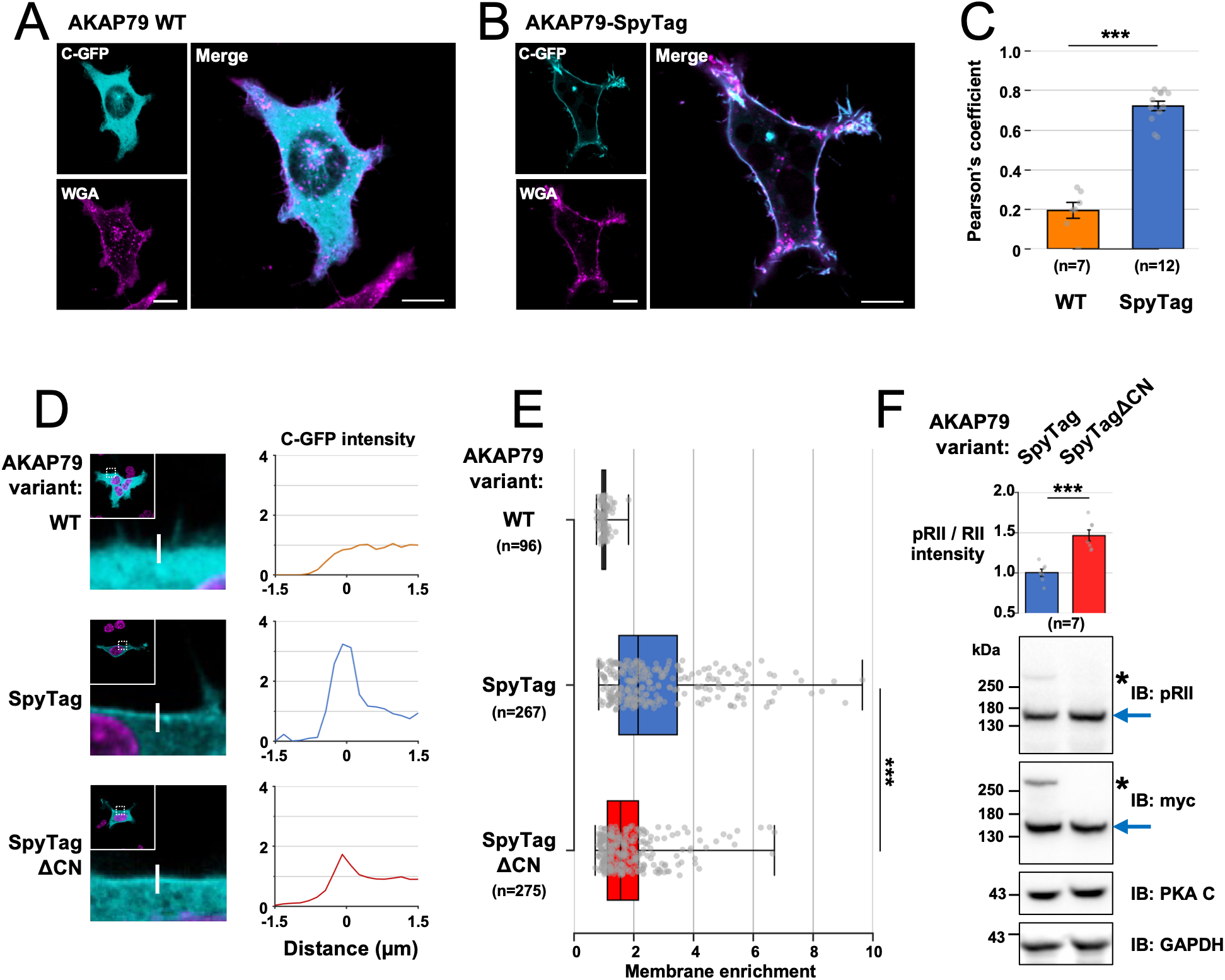
Imaging C subunit membrane enrichment driven by AKAP79 anchoring. Confocal images of SpyC-RII cells transfected with C-GFP and either the WT (**A**) or SpyTag (**B**) variant of AKAP79. Cells were imaged live for C-GFP (cyan) after staining with WGA-CF633 (magenta). Scale bars correspond to 10 µm. (**C**) Quantification of C-GFP/WGA-CF633 colocalization. (**D**) Example line scans of SpyC-RII cells transfected with C-GFP and either WT (upper sub-panel), SpyTag (middle) or SpyTagΔCN (bottom) variants of AKAP79. Live images are shown in the lefthand column with the C-GFP channel in cyan, and Hoechst-33342 stain in magenta. Line intensities are normalized to the peak value. (**E**) Box plots showing distributions of membrane enrichment (intensity at peak/1.5 µm inside cell) for the conditions shown in the preceding panel. Medians (straight grey lines), and 25^th^/75^th^ percentiles (dotted lines) are shown. (**F**) Comparison of RII phosphorylation by IB densitometry using protein extracts from cells transfected with either SpyTag or SpyTagΔCN AKAP79. For panels C and F, statistical comparisons were performed using unpaired two-tailed Student’s t-tests (\*\*\**P* < 0.001).

Previous studies show that AKAP79/150-associated calcineurin can exert tonic control over local signalling in unstimulated cells^33,34^. We reasoned that if CN does increase the proportion of dephosphorylated RII subunits bound to AKAP79 to a meaningful degree, this should elevate the rate of C subunit capture within the complex with detectable effects on membrane enrichment in our system. We therefore investigated the effects of removing the CN anchoring motif between positions 337-343 (‘ΔCN’) of AKAP79 on C subunit membrane enrichment. We measured C subunit membrane enrichment using cell edge line scanning of SpyC-RII cells co-expressing C-GFP with different AKAP79 variants (**Fig. 3D**). For each cell, we calculated a membrane enrichment score by dividing peak C-GFP intensity at the cell edge by the intensity 1.5 μm into the cytosol. SpyC-RII cells expressing AKAP79-WT exhibited no membrane enrichment of C subunits as expected (ratio of 1.01 ± 0.02, n = 96, **Fig. 3D**, upper panel), whereas C subunits were markedly enriched at the membrane of cells expressing AKAP79-SpyTag (2.75 ± 0.1, n = 267, middle panel). Membrane enrichment was reduced by about half in cells expressing AKAP79ΔCN-SpyTag (1.85 ± 0.06, n = 275) according to mean enrichment ratios (**Fig. 3E**). The unique mobility of different AKAP-RII conjugates using the SpyC-RII systems also enables phosphorylation levels of RII bound to specific AKAPs to be measured. We took advantage of this feature to calculate pRII/RII ratios in extracts from SpyC-RII cells expressing either AKAP79-SpyTag or AKAP79-SpyTag ΔCN (**Fig 3F**) using IB densitometry. The fraction of pRII subunits was ∼ 46 % higher in extracts from cells expressing AKAP79-SpyTagΔCN (normalized pRII/RII ratio of 1.46 ± 0.07) compared to AKAP79-SpyTag (1.0 ± 0.04). This confirms that the proportion of dephosphorylated RII subunits is higher in the AKAP79 complex when the CN anchoring site is present. In sum, these experiments confirm an important role for phosphatase co-anchoring in regulating C subunit dynamics, and highlight the novel experimental possibilities afforded by our isopeptide-based selective anchoring system.

### RII anchoring to MAP2 supports microtubule straightening

Although AKAP79 is a well-established organizer of signaling in dendritic spines, the major PKA-anchoring protein in dendritic shafts is MAP2. MAP2 was the first AKAP to be identified^35^ and it is the predominant determinant of PKA localization in neurons^36^. Genetic and imaging studies support a role for MAP2 in dendritic stability and microtubule organization: MAP2 deletion reduces dendritic length and microtubule density and eliminates prominent microtubule cross-bridges visible by deep-etch electron microscopy^37^. However, the functional consequences of PKA anchoring to MAP2 are not fully resolved. Native MAP2 – with associated interaction partners – enhances microtubule bundling *in vitro*, whereas MAP2 stripped of its interaction partners does not^38^, hinting at a role for RII in stabilizing microtubule bundles. Conversely, biochemical studies have shown that PKA phosphorylation of MAP2 reduces its affinity for microtubules^39,40^, raising the possibility that PKA anchored at MAP2 might rather destabilize microtubules. We aimed to use our selective anchoring system to clarify this uncertainty by isolating the contribution of RII anchored to MAP2. For all experiments, we focused on a high-molecular-weight (HMW) MAP2 sequence, as HMW MAP2 splice variants predominate in mature neurons, including mature hippocampal neurons^40^. We first validated SpyTag-mediated RII anchoring to MAP2 by live imaging SpyC-RII cells co-transfected with C-GFP and either the WT or SpyTag variant of MAP2. We imaged both C-GFP subunits (cyan, **Fig. 4A–B**) and microtubules stained with Tubulin Tracker Deep Red (magenta, **Fig. 4A–B**). Whereas C-GFP remained diffusely distributed throughout the cytosol in cells expressing MAP2-WT (**Fig. 4A**), it was clearly sequestered to microtubules when MAP2-SpyTag was present (**Fig. 4B**). Quantification using Pearson’s correlation coefficient confirmed robust colocalization of C-GFP with microtubules only in cells expressing MAP2-SpyTag (MAP2-WT: 0.22 ± 0.03; MAP2-SpyTag: 0.85 ± 0.02; *P* < 0.001; **Fig. 4C**).

**Figure 4.**
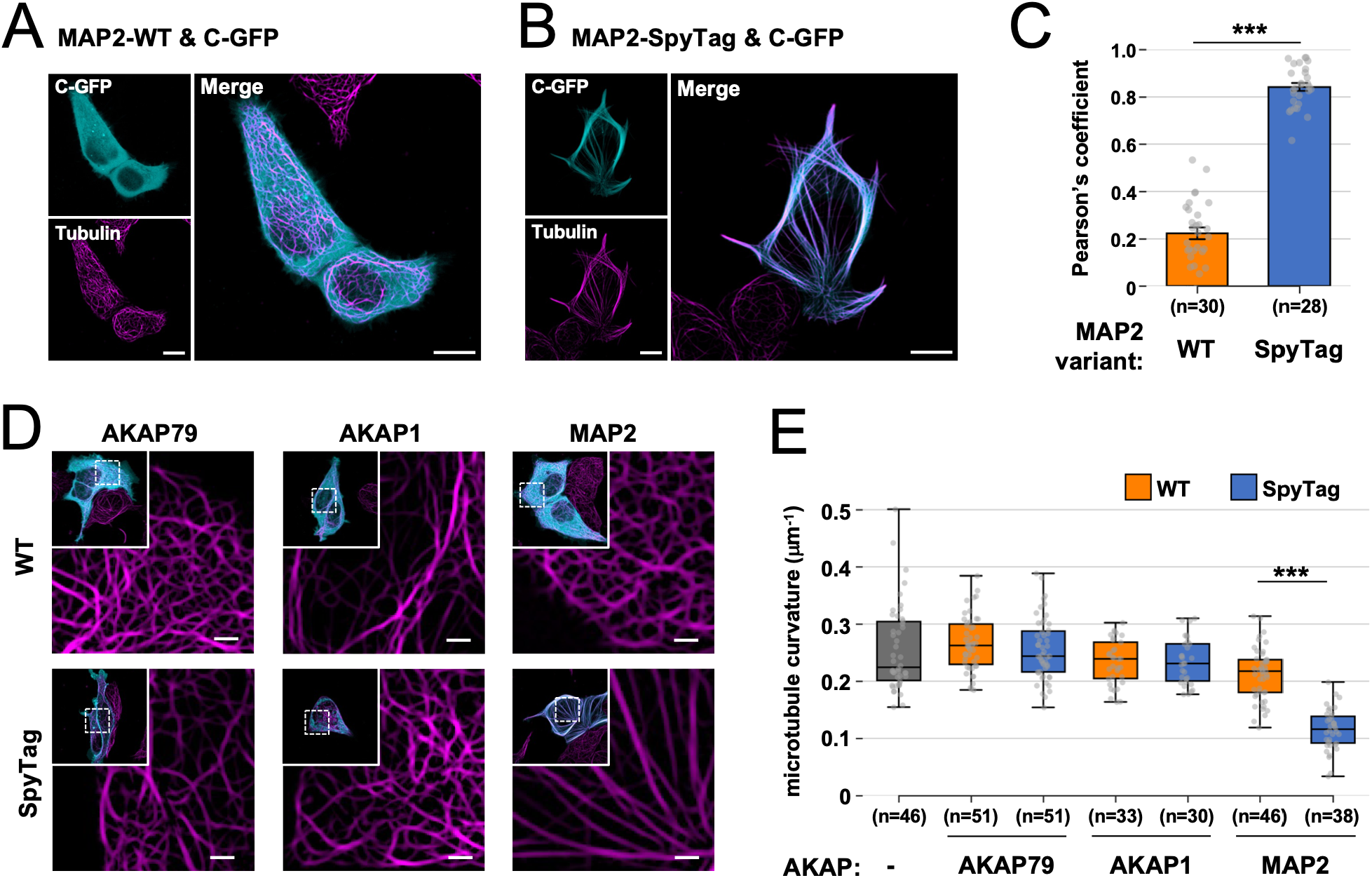
MAP2-bound RII induces microtubule straightening. Confocal images of live SpyC-RII cells co-transfected with C-GFP and either the WT (**A**) or SpyTag (**B**) variant of MAP2. Cells were stained with Tubulin Tracker Deep Red (magenta), which is shown alongside C-GFP signal (cyan). Scale bars correspond to 10 µm. (**C**) Quantification of C-GFP/Tubulin Tracker colocalization for the conditions shown in the preceding panels. (**D**) Comparison of microtubule structure for cells co-transfected with C-GFP and either the WT (upper row) or SpyTag (lower row) variants of AKAP79, AKAP1, and MAP2. Cells were stained with Tubulin Tracker Deep Red (magenta), and only this channel is shown for clarity. Scale bars correspond to 2 µm. (**E**) Quantification of microtubule curvature in SpyC-RII cells transfected with empty vector (grey), or according to the three WT (orange) and SpyTag (blue) AKAP conditions shown in the preceding panel. Statistical comparison of colocalisation data (panel C) was performed using a two-tailed unpaired Student’s t-test. Differences in microtubule curvature between WT and SpyTag pairs were assessed using two-tailed Mann Whitney U tests with Bonferroni correction for multiple comparisons (\*\*\**P* < 0.001).

While confirming that SpyTag-mediated anchoring efficiently recruits SpyC-RII to MAP2-positive microtubules, we also noted a striking change in microtubule organization in cells expressing MAP2-SpyTag. In addition to the robust redistribution of C-GFP onto microtubules, the microtubule network appeared markedly straighter and more aligned in this specific condition (**Fig. 4B**). We therefore quantified microtubule curvature across cells expressing WT and SpyTag variants of MAP2, AKAP79 and AKAP1. Microtubules appeared straighter in SpyC-RII cells co-expressing C-GFP with MAP2-SpyTag than with MAP2-WT (right column, **Fig 4D**). This straightening effect was not observed in SpyC-RII cells expressing either variant of AKAP79 (left column, **Fig. 4D**) or AKAP1 (middle column). We quantified microtubule curvature by fitting to B-spline curves: smaller μm^-1^ values indicate straighter filaments according to this approach^41^. Quantitative analysis confirmed a marked reduction in microtubule curvature in cells expressing MAP2-SpyTag (0.12 ± 0.01 μm⁻¹) compared with MAP2-WT (0.21 ± 0.01 μm⁻¹; *P* < 0.001; **Fig. 4E**). Mean microtubule curvature was slightly reduced in MAP2-WT cells compared to all other conditions (control, AKAP79-WT & AKAP79-SpyTag: 0.26 ± 0.01 μm^-1^; AKAP1-WT & AKAP1-SpyTag: 0.23 ± 0.01 μm^-1^, **Fig. 4E**). Taken together, these data indicate that recruitment of RII to MAP2 markedly straightens microtubules, consistent with a specific effect of the MAP2–RII complex on microtubule organization.

### The effects of AKAP1 but not MAP2 anchoring depend on PKA activity

A conventional interpretation of functional effects linked to a specific PKA anchoring site is that the anchoring protein promotes local phosphorylation of nearby substrates whose phosphorylated state mediates the downstream effect. However, in the case of MAP2, such an interpretation is at odds with previous reports that PKA phosphorylates MAP2 to trigger its release from microtubules^40^. Given the very high concentrations of both MAP2 and RII in dendritic shafts^13,36^ and the growing recognition that abundant neuronal signaling enzymes such as CaMKII and SynGAP can also serve important structural functions^16,17^, we considered the alternative possibility that RII bound to MAP2 contributes structurally to microtubule organization. Before testing whether the MAP2–RII microtubule-straightening effect depended on C activity, we first asked whether our system could recapitulate a known phosphorylation-dependent anchoring effect using AKAP1.

AKAP1 is a scaffold protein that recruits PKA to the outer mitochondrial membrane (OMM)^42,43^ for local PKA phosphorylation of Drp1 that suppresses mitochondrial fission^44,45^. AKAP1 also anchors phosphatases CN and PP1^46^, which oppose PKA at the OMM^47^. Consistent with this tightly balanced phospho-signaling system, basal PKA phosphorylation is elevated at the OMM relative to other compartments and is rapidly lost following PKA inhibition with H-89^48^. We therefore reasoned that expression of AKAP1-SpyTag in SpyC-RII cells should recruit C-subunits to mitochondria and provide a suitable benchmark for a catalytic, phosphorylation-dependent anchoring effect. To test this, we co-expressed C-GFP with either control empty vector, AKAP1-WT or AKAP1-SpyTag in SpyC-RII cells and compared average mitochondrial size across these conditions using live imaging following staining with MitoView-633 (magenta, **Fig. 5A**). Mitochondria in control cells exhibited a mean area of 0.524 +/- 0.032 µm^2^, which fell to 0.248 +/-0.011 µm^2^ when AKAP1-WT was included (**Fig. 5A–B**), consistent with unopposed phosphatase activity favoring mitochondrial fission in the absence of recruited PKA activity. Inclusion of AKAP1-SpyTag returned mitochondrial size to similar levels to the control (0.674 +/- 0.062 µm^2^) with the distribution including a few cells exhibiting very large disc shaped mitochondria lined by C-GFP subunits (**Fig. 5A**, lower right sub-panel). We next tested the sensitivity of this effect to PKA C subunit inhibition with 10 μM H-89. Whereas H-89 had no detectable effect on cells expressing AKAP1-WT (vehicle: 0.216 ± 0.004 µm^2^, H-89: 0.224 ± 0.008 µm^2^), mitochondrial area was markedly reduced for AKAP1-SpyTag-expressing cells (vehicle: 0.602 ± 0.046 µm^2^, H-89: 0.355 ± 0.034 µm^2^, *P* < 0.001, **Fig. 5C–D**). Having confirmed that our selective anchoring system can recapitulate a bona fide phosphorylation-dependent AKAP phenotype, we next asked whether the MAP2–RII microtubule-straightening effect showed the same sensitivity to H-89.

**Figure 5.**
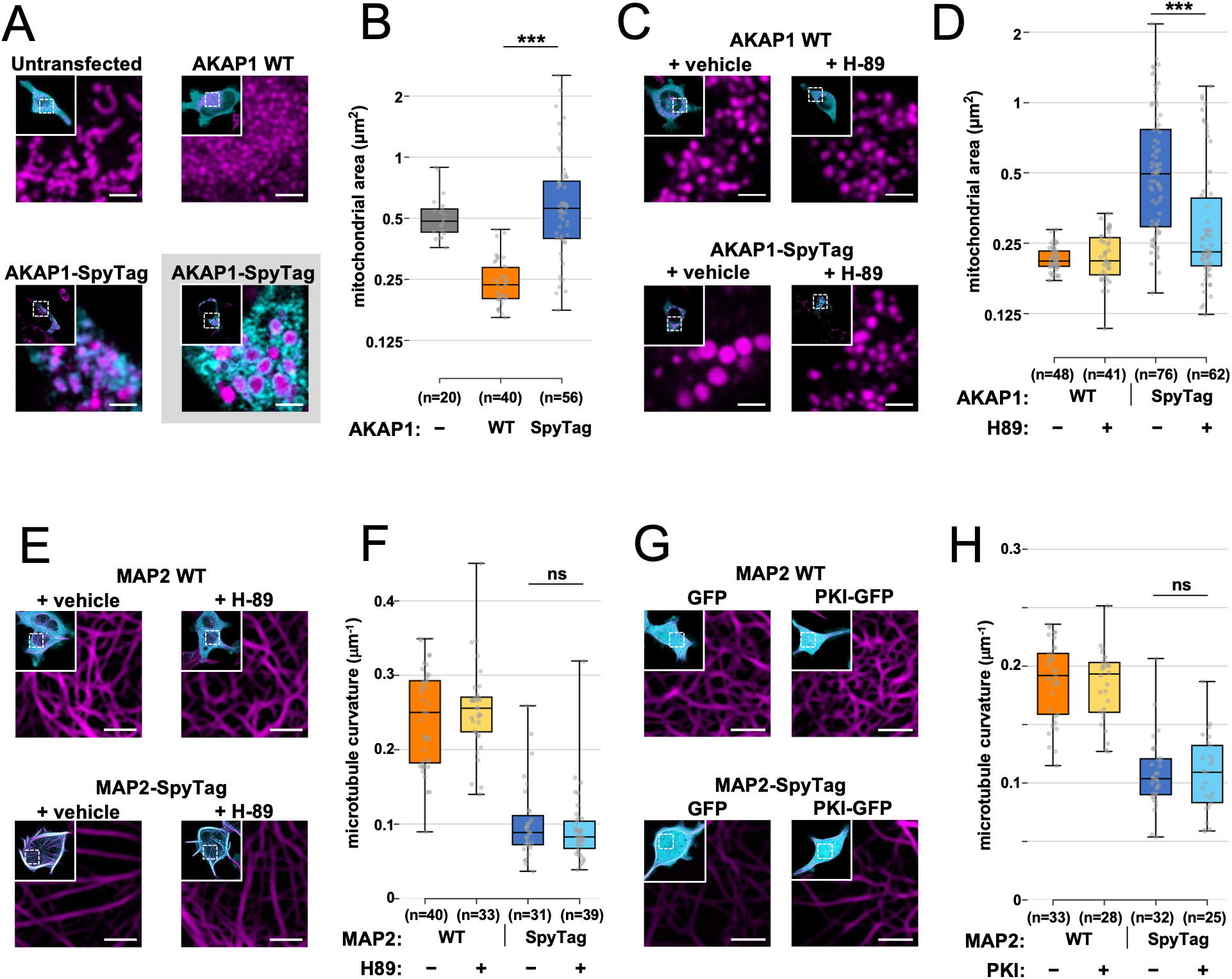
PKA activity dependence of AKAP1 and MAP2 anchoring functions. Panels **A**, **C**, **E** and **G** show live-cell confocal images of SpyC-RII cells under the indicated conditions (scale bars, 10 µm). In **A** and **C**, mitochondria stained with MitoView-633 are shown in magenta. In **E** and **G**, microtubules stained with Tubulin Tracker are shown in magenta. (**A**) SpyC-RII cells co-transfected with C-GFP and either empty vector or WT/SpyTag AKAP1. The lower right sub-panel shows enlarged mitochondria lined with C subunits observed in a subset of AKAP1-SpyTag cells (C-GFP, cyan). (**B**) Boxplot of mean area of individual mitochondria within each cell for the conditions in A (log2 scale). (**C**) SpyC-RII cells transfected with C-GFP and either WT (top) or SpyTag (bottom) AKAP1 following 24 h pre-treatment with vehicle or 10 µM H-89. (**D**) Boxplot of mitochondrial area distributions for the conditions in C. (**E**) SpyC-RII cells co-transfected with C-GFP and either WT (top) or SpyTag (bottom) MAP2 following 24 h pre-treatment with vehicle or 10 µM H-89. (**F**) Quantification of microtubule curvature for the conditions in E. (**G**) SpyC-RII cells co-transfected with MAP2 WT (top) or MAP2-SpyTag (bottom), together with either GFP (left) or PKI-GFP (right). (**H**) Quantification of microtubule curvature for the conditions shown in **G**. For all boxplots, boxes indicate the interquartile range (25th–75th percentiles), the centre line marks the median, and whiskers show the full data range. Statistical comparisons were performed using two-tailed Mann–Whitney tests (\*\*\**P* < 0.001; ns, *P* > 0.5).

To test whether MAP2-dependent microtubule straightening requires PKA catalytic activity, SpyC-RII cells co-expressing C-GFP with either MAP2-WT or MAP2-SpyTag were pre-incubated with the PKA inhibitor H-89 (10 µM) or vehicle (DMSO) for 24 h prior to imaging, with treatment maintained throughout image acquisition. In vehicle-treated cells, MAP2-SpyTag expression induced robust microtubule straightening relative to MAP2-WT (MAP2-WT: 0.24 ± 0.01 μm^-1^; MAP2-SpyTag: 0.10 ± 0.01 μm^-1^; p < 0.001; **Fig. 5E–F**). In contrast to the AKAP1 anchoring phenotype (**Fig. 5C–D**), inhibition of PKA catalytic activity with H-89 did not attenuate microtubule straightening in MAP2-SpyTag-expressing cells, with curvature values indistinguishable from vehicle controls (vehicle: 0.10 ± 0.01 μm^-1^; H-89: 0.09 ± 0.01 μm^-1^; p = 0.51; **Fig. 5E–F**). To further assess whether microtubule straightening persists under sustained suppression of PKA activity, we co-expressed MAP2-WT or MAP2-SpyTag with either GFP or the peptide PKA inhibitor PKI fused to GFP (PKI-GFP), which inhibits PKA-C subunits^49^. Consistent with the H-89 experiments, MAP2-SpyTag promoted microtubule straightening relative to MAP2-WT when co-expressed with GFP alone (MAP2-WT: 0.19 ± 0.01 μm^-1^; MAP2-SpyTag: 0.11 ± 0.02 μm^-1^; p < 0.001; **Fig. 5G–H**). Expression of PKI-GFP did not prevent MAP2-SpyTag-dependent straightening across three independent experiments (MAP2-SpyTag + PKI-GFP: 0.11 ± 0.03 μm^-1^), with no difference relative to GFP controls (p = 0.94; **Fig. 5H**). Together, these data indicate that MAP2-associated microtubule straightening does not require PKA catalytic activity and is instead consistent with a phosphorylation-independent role for RII anchoring along microtubules.

### Mechanistic insights into the structural role of RII subunits anchored to MAP2

We next examined isoform dependence and began to define the molecular basis of the microtubule straightening effect, while corroborating our observations in a system that did not rely on isopeptide-based anchoring. To this end, we performed experiments in ΔRII HEK293T cells, expressing RII and MAP2 constructs with intact native anchoring interfaces. Microtubule curvature was compared in ΔRII cells co-expressing MAP2 with either empty vector, RIIα, or RIIβ. Co-expression of either RII isoform induced significant microtubule straightening, with mean curvature values of 0.11 ± 0.01 µm⁻¹ for RIIα and 0.12 ± 0.01 µm⁻¹ for RIIβ, compared to 0.18 ± 0.01 µm⁻¹ for MAP2 alone (**Fig. 6A–B**). To further delineate the structural requirements for this effect, we compared microtubule curvature in cells expressing MAP2 together with either full-length RIIα or RIIα truncation mutants lacking the CNB-B domain (ΔB) or both CNB domains (ΔAB) (**Fig. 6C**). Whereas full-length RIIα produced robust microtubule straightening (mean curvature 0.09 ± 0.01 µm^-1^, *P* < 0.001), curvature values when both CNB domains were absent were indistinguishable from the control condition (0.21 ± 0.01 µm⁻¹ with no RII; 0.21 ± 0.01 µm⁻¹ with RIIα ΔAB). RIIα lacking only CNB-B supported only a limited straightening effect at most (0.18 ± 0.01 µm⁻¹) (**Fig. 6D–E**). These experiments confirm that the microtubule straightening effect of RII anchored to MAP2 extends to the native anchoring mode and is consistent with an essential role for the second CNB domain in this process.

**Figure 6.**
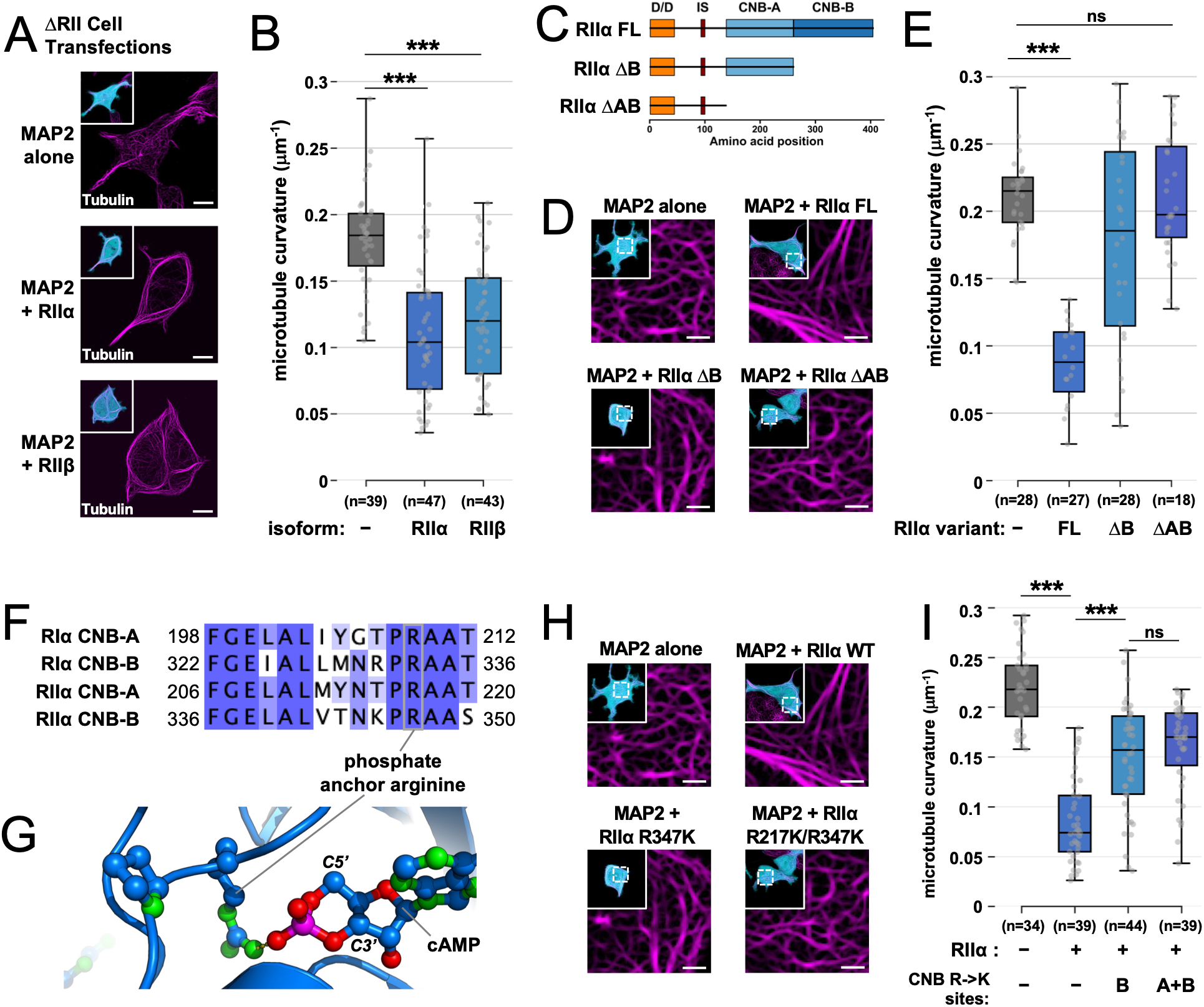
Molecular determinants of microtubule straightening by MAP2-anchored RII. Panels **A**, **D** and **H** show confocal images of live ΔRII cells co-transfected with GFP and the proteins indicated. GFP signal (cyan) is shown in the inset images, with Tubulin Tracker Deep Red stain shown in magenta. Scale bars correspond to 10 µm (**A**) and 2 µm (**D**, **H**). Panels **B**, **E** & **I** show quantification of microtubule curvature for the conditions shown in the **A**, **D**, and **H**, respectively. (**C**) Domain topology of RIIα truncation constructs. (**F**) Alignment of phosphate binding cassettes of human RIα and RIIα. (**G**) Structure showing coordination of the 3’-5’ cyclic phosphate by the phosphate-anchor arginine in RIIα (PDB 4JVA). For all boxplots, boxes indicate the interquartile range (25th–75th percentiles), the centre line marks the median, and whiskers show the full data range. Statistical comparisons were performed using two-tailed Mann–Whitney tests; \*\*\**P* < 0.001, ns, *P* > 0.5.

Each PKA regulatory subunit CNB domain contains a phosphate-binding cassette with an invariant arginine anchor residue that coordinates the cyclic phosphate of cAMP (**Fig. 6F–G**)^50,51^. Previous studies of native MAP2 preparations showed that MAP2-associated binding partners increase microtubule solution viscosity, and that addition of cAMP enhances this effect further^37^. Together with our truncation data, this suggested that cAMP binding to MAP2-associated RII may contribute to the full microtubule straightening phenotype. We therefore introduced arginine-to-lysine substitutions into the RIIα phosphate-binding cassette, targeting either the higher-affinity CNB-B site alone (R347K) or both CNB-A and CNB-B sites (R217K/R347K). Whereas wild-type RIIα strongly reduced microtubule curvature from 0.22 ± 0.01 µm⁻¹ to 0.085 ± 0.01 µm⁻¹, mutation of CNB-B produced only partial straightening (0.15 ± 0.01 µm⁻¹, *P* < 0.001 versus RIIα WT) at comparable levels to when both sites were mutated (0.16 ± 0.01 µm⁻¹; ns versus R347K only; **Fig. 6H–I**). These data suggest that the integrity of the CNB-B cAMP-binding pocket is required for maximal MAP2-RII-dependent microtubule straightening, consistent with earlier viscosity measurements in which native MAP2 complexes increased microtubule organization and cAMP further enhanced this effect.

The microtubule straightening phenotype, together with previous studies linking MAP2 to increased solution viscosity^38^ and microtubule cross-linking^37^, suggested that RII anchored to MAP2 might influence microtubule bundling. To test this directly, we used correlative light and electron microscopy (CLEM) to identify SpyC-RII cells co-expressing C-GFP with either MAP2-WT or MAP2-SpyTag, and subsequently examined microtubule organization by transmission electron microscopy of 70 nm sections (**Fig. 7A, D**). Additional examples without microtubule annotations are shown in **figure S3**. Previous ultrastructural studies have shown that MAP2 can promote formation of microtubule bundles characterized by unusually wide inter-filament spacing^40^. Consistent with the absence of RII recruitment, microtubules in MAP2-WT cells were typically observed as isolated filaments or as small bundles containing only a few closely packed microtubules (**Fig. 7A**). Quantification confirmed that most structures contained one to three filaments, with a median bundle size of two filaments (**Fig. 7B**). Where parallel microtubules were observed, centre-to-centre spacing was typically 30–40 nm (median ∼35 nm; **Fig. 7C**).

**Figure 7.**
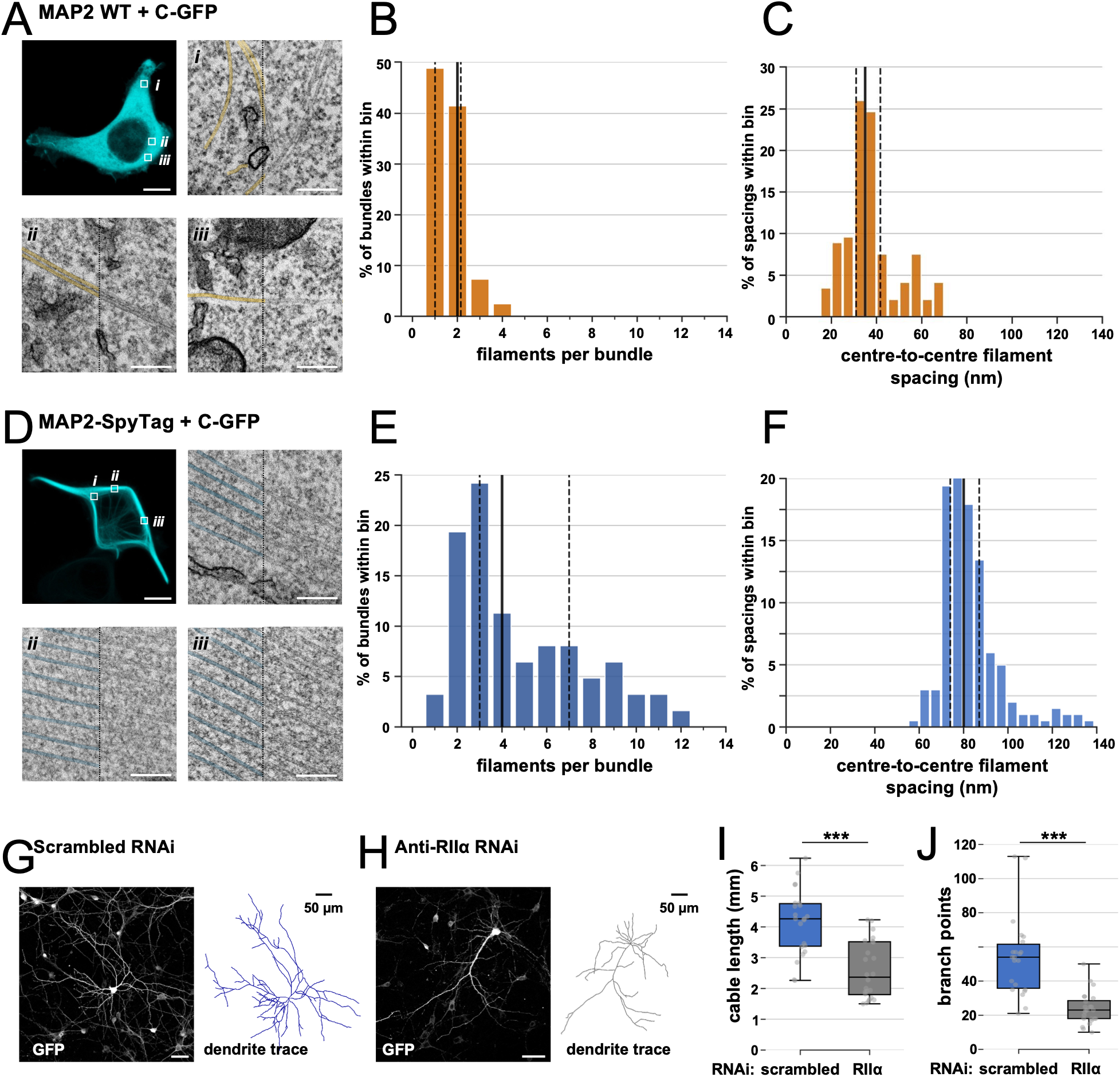
MAP2-anchored RII promotes microtubule bundling and supports dendritic arborization. (**A**, **D**) CLEM of SpyC-RII cells co-expressing C-GFP with either MAP2-WT (**A**) or MAP2-SpyTag (**D**). Fluorescence imaging was used to identify transfected cells and regions of interest (upper left sub-panels). Representative electron micrographs from the indicated regions are shown to the right and below. Scale bars: fluorescence images, 10 µm; electron micrographs, 200 nm. Microtubule filaments are highlighted on the left-hand half of micrographs in orange (**A**) and cyan (**D**). (**B**, **E**) Distribution of microtubule bundle sizes quantified from electron micrographs of cells expressing MAP2-WT (**B**; 82 bundles from 3 cells) or MAP2-SpyTag (**E**; 62 bundles from 3 cells). Histograms show the percentage of bundles containing the indicated number of parallel microtubule filaments. (**C**, **F**) Distribution of centre-to-centre spacing distances between neighbouring parallel microtubule filaments within bundles from MAP2-WT (**C**; 146 measurements from 3 cells) and MAP2-SpyTag (**F**; 208 measurements from 3 cells). Histograms show the percentage of measurements within each bin. In panels **B**, **C**, **E** and **F**, dashed lines indicate the 25th and 75th percentiles and the solid line indicates the median. (**G**, **H**) Representative images of cultured hippocampal neurons expressing either scrambled control RNAi (**G**) or RNAi targeting RIIα (**H**). GFP fluorescence images are shown alongside dendritic tracings used for morphometric analysis. (**I**, **J**) Quantification of total dendritic cable length per neuron (**I**) and total branch points per neuron (**J**) following expression of scrambled control RNAi or RIIα-targeting RNAi. Box plots show the median, interquartile range and minimum-to-maximum values. Statistical comparisons were performed using two-tailed Mann-Whitney U tests. \*\*\**P* < 0.001.

In contrast, MAP2-SpyTag cells displayed prominent arrays of parallel microtubules extending across large regions of cytoplasm (**Fig. 7D**). These arrays frequently contained substantially larger numbers of aligned filaments than those observed in MAP2-WT cells (**Fig. 7E**), with a median bundle size of five filaments. Notably, bundled microtubules in MAP2-SpyTag cells exhibited markedly larger inter-filament spacing than in MAP2-WT cells. Distances clustered around a median centre-to-centre separation of ∼80 nm (**Fig. 7F**). These dimensions closely resemble the unusually wide spacing reported for MAP2-associated microtubule bundles in dendrites ^40^. Finally, to test whether RIIα contributes to neuronal morphology, we depleted RIIα in primary hippocampal neurons, the predominant dendritic RII isoform in these cells. RIIα knockdown reduced total dendritic cable length by approximately 38%, from 4.1 ± 0.2 mm to 2.6 ± 0.2 mm (*P* < 0.001; **Fig. 7I**), and reduced branch point number from 54 ± 5 to 24 ± 2 per neuron (*P* < 0.001; **Fig. 7J**). This reduction in both dendrite length and branching suggests that RIIα may contribute not only to dendritic extension but also to the stabilization or maintenance of newly formed dendritic branches. Together, these ultrastructural and neuronal data support a structural role for MAP2-anchored RII in promoting widely spaced microtubule bundles and maintaining dendritic arborization.

## Discussion

Compartmentalized cAMP signaling is organized by AKAPs, yet the function of individual anchored PKA pools has remained difficult to resolve because most perturbations do not isolate individual anchoring sites. Here, we addressed that limitation by engineering a replacement system in which endogenous type II PKA regulatory subunits are exchanged for a SpyCatcher-tagged variant that is selectively re-anchored only to AKAPs bearing a SpyTag-substituted docking site. This strategy allowed us to compare AKAP effects with and without site-specific PKA anchoring in living cells while leaving the broader signaling environment largely intact. Using this platform, we validated selective reconstitution of compartment-specific PKA behavior in the AKAP79 and AKAP1 complexes, confirming that CN action within the AKAP79 complex draws PKA C subunits into the complex (**Fig. 3**), and that AKAP1 directs PKA to the OMM for regulation of mitochondrial fusion (**Fig. 5**). The principal insight, however, came from investigation of MAP2: Anchoring of RII to MAP2 drove a marked microtubule-straightening phenotype that persisted even with pharmacological or peptide-mediated inhibition of PKA catalytic activity and was recapitulated using native MAP2–RII binding rather than isopeptide conjugation. Together, these findings support a model in which anchored RII subunits can fulfil either structural or enzymatic roles, depending on the specific AKAP complex. Thus, the present work provides a generalizable tool for interrogating AKAP-specific signaling and challenges the assumption that the biological effects of anchored PKA are necessarily mediated by local phosphorylation.

Measurements of PKA C subunit association with AKAP79 build on our previous work showing that pRII dephosphorylation by CN within the AKAP79 complex reduces PKA activity^14^. Our measurements in SpyC-RII cells (**Fig. 3**) support that role while revealing an additional potential function for phosphatase co-anchoring, namely the selective regeneration of PKA holoenzymes at particular subcellular sites. In this framework, phosphatases can sharpen compartment identity by biasing where liberated C subunits are recaptured. Membranes may be particularly favorable environments for such a mechanism, given that the N-myristoyl groups of PKA C subunits are thought to embed in the cell membrane upon release from RII subunits, which would be expected to further increase the likelihood of local rebinding^52,53^. More broadly, these findings imply that different anchored pools of RII engage in a ‘tug-of-war’ for a limited pool of C subunits, with local dephosphorylation of RII subunits and local cAMP signals dictating the distribution of the C subunits^11,14^. In neurons, where Ca²⁺ influx and neuromodulator-driven cAMP signals overlap in both time and space, this logic may be particularly important with the output of an AKAP complex depending as much on its ability to retain PKA C subunits as its ability to release them at the right time and place.

MAP2 is among the most abundant neuronal proteins and the dominant dendritic PKA-anchoring protein, and it has long been implicated in microtubule spacing, bundling, and stabilization^36,38,54^. Conversely, previous work with purified microtubules has shown that PKA phosphorylation of MAP2 leads to release of MAP2 from microtubules^39,40^, which would be expected to have the opposite effects. Our findings help to reconcile these observations. The persistence of microtubule straightening under both H-89 and PKI inhibition shows that the anchored regulatory subunit itself contributes to microtubule organization independently of PKA catalytic activity. This interpretation is strengthened by the finding that both RII isoforms support microtubule straightening, whereas loss of the second CNB domain, or mutation of its cAMP-binding pocket, markedly reduces the phenotype. Importantly, our EM analysis extends this conclusion beyond filament straightening by showing that MAP2-anchored RII promotes the formation of larger, widely spaced microtubule bundles. In MAP2-WT cells, microtubules were typically isolated or present in small, closely packed bundles, whereas MAP2-SpyTag cells contained extended arrays of parallel filaments with increased bundle size and a shift in centre-to-centre spacing from approximately 35 nm to approximately 80 nm. These ultrastructural changes closely resemble the wide spacing associated with MAP2-containing dendritic microtubule bundles and suggest that RII stabilizes or remodels MAP2-dependent crosslinking rather than simply altering microtubule curvature. More broadly, these data place anchored PKA alongside an increasing number of abundant neuronal signaling proteins, including CaMKII and SynGAP, whose organizational roles have emerged only after years of being viewed primarily through a catalytic lens^16,17^.

Earlier studies disrupting the MAP2 PKA-binding region or deleting MAP2 entirely revealed defects in dendritic architecture, contextual learning, and norepinephrine-enabled plasticity, which have generally been interpreted as consequences of impaired local phosphorylation^36,37,40,55^. Our results do not argue against phosphorylation-based mechanisms, but they suggest that at least part of the MAP2-PKA phenotype arises from loss of a structural MAP2-RII assembly. This interpretation is supported by the formation of large, widely spaced microtubule bundles when RII is recruited to MAP2, by the reduction in dendrite cable length and branch point number following RIIα depletion, and by the requirement for the CNB-B domain and its cAMP-binding pocket for maximal microtubule straightening. These findings also resonate with studies showing that cAMP signaling promotes neuronal maturation and dendritic arborization, including increased branch number and dendritic length following PDE4 inhibition or cAMP analogue treatment in hippocampal neurons^56^, and with evidence that cAMP gradients in developing neurons are shaped by AKAP-anchored feedback mechanisms^57^. Together with earlier observations that native MAP2 complexes increase microtubule viscosity and that cAMP further enhances this effect, our data raise the possibility that MAP2-RII complexes respond to cAMP across distinct concentration regimes. At very low cAMP, RII may remain largely unliganded, with little effect on either microtubule organization or C-subunit release. At low or basal cAMP, preferential occupancy of the higher-affinity CNB-B site could promote a structural state of RII that enhances MAP2-dependent microtubule bundling without triggering substantial C-subunit release. At higher cAMP, occupancy of both CNB domains would be expected to drive canonical PKA activation, adding local phosphorylation to this structural mode of regulation. In this model, neuromodulator-driven cAMP signals could influence dendritic architecture not only by regulating transcriptional or phosphorylation-dependent maturation pathways, but also by acutely changing the composition, conformation or mechanics of cytoskeletal scaffolds. Future work should define the molecular interface between MAP2, RII and microtubules, determine how cAMP binding and C-subunit occupancy alter this assembly, and establish how MAP2-RII-dependent bundling affects dendritic trafficking, branch stabilization, spine plasticity and electrical integration. In that sense, anchored PKA complexes should be viewed not only as spatially restricted signaling enzymes, but also, in some contexts, as regulated structural components of the cell.

### Limitations of the study

This study does not yet define the molecular interface through which MAP2-bound RII alters microtubule organization, or whether RII contacts microtubules directly or stabilizes MAP2-dependent crosslinks. Although cAMP-binding mutants implicate the CNB-B cAMP-binding pocket in maximal microtubule straightening, we did not directly manipulate local cAMP concentrations or measure RII nucleotide occupancy in the MAP2 complex. Finally, RIIα depletion reduced dendrite length and branch point number in hippocampal neurons, but these experiments do not distinguish effects on branch initiation, growth, stabilization or survival, nor consider potential effects on spine structure^58^. Future structural, biophysical and neuronal imaging approaches will be needed to define how cAMP binding, C-subunit occupancy and MAP2-RII assembly regulate dendritic microtubule architecture.

## STAR★Methods

## Key resource table

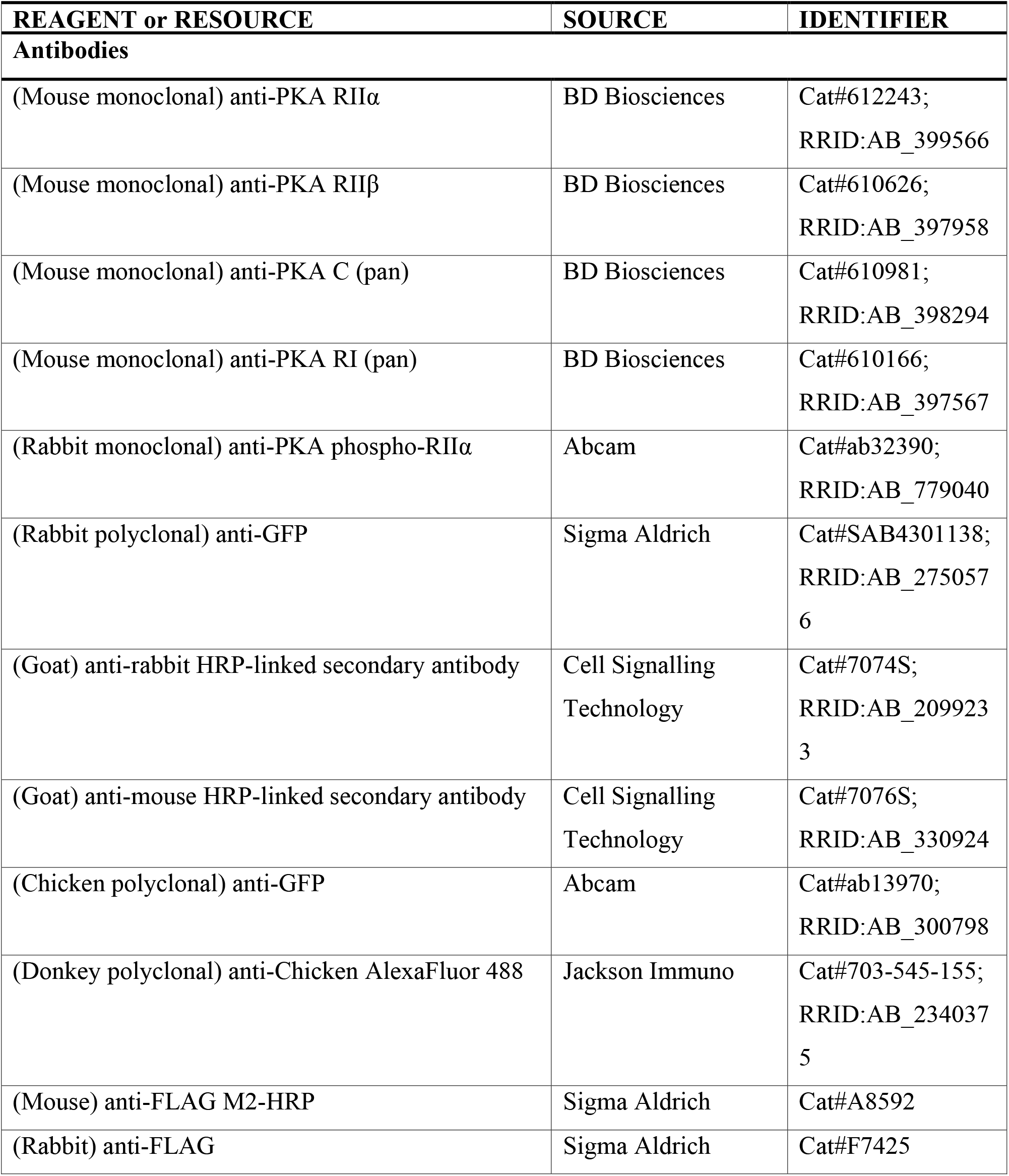
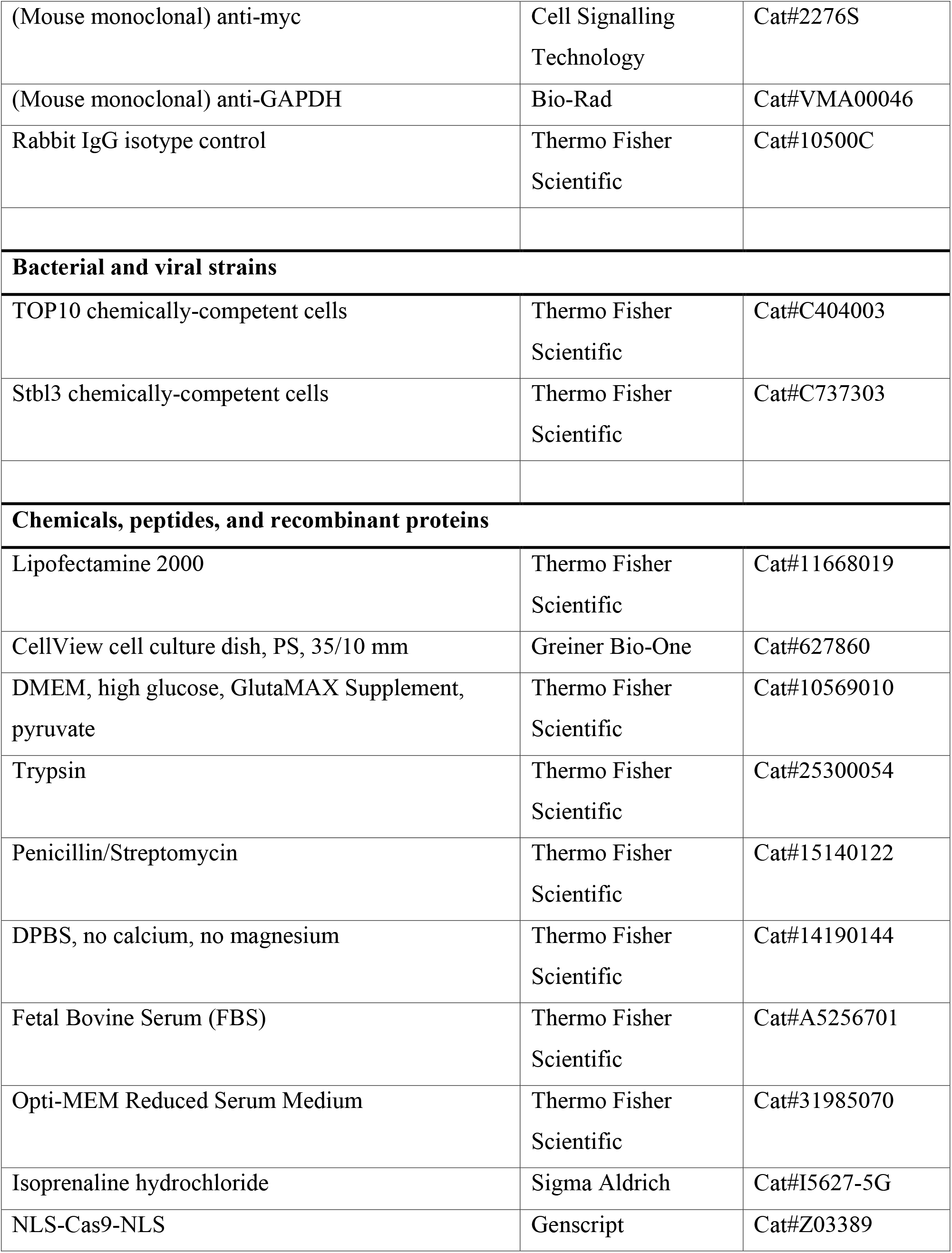
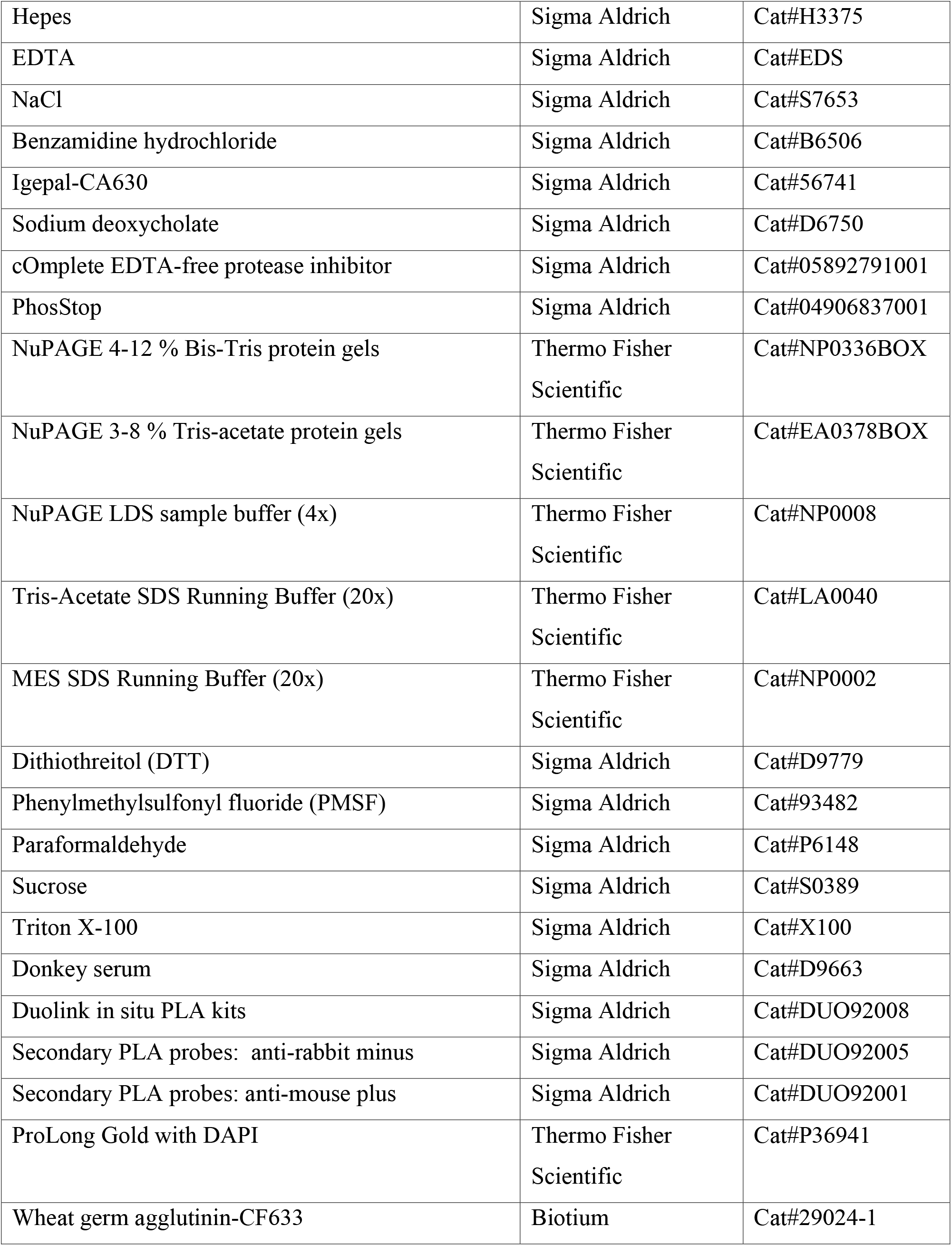
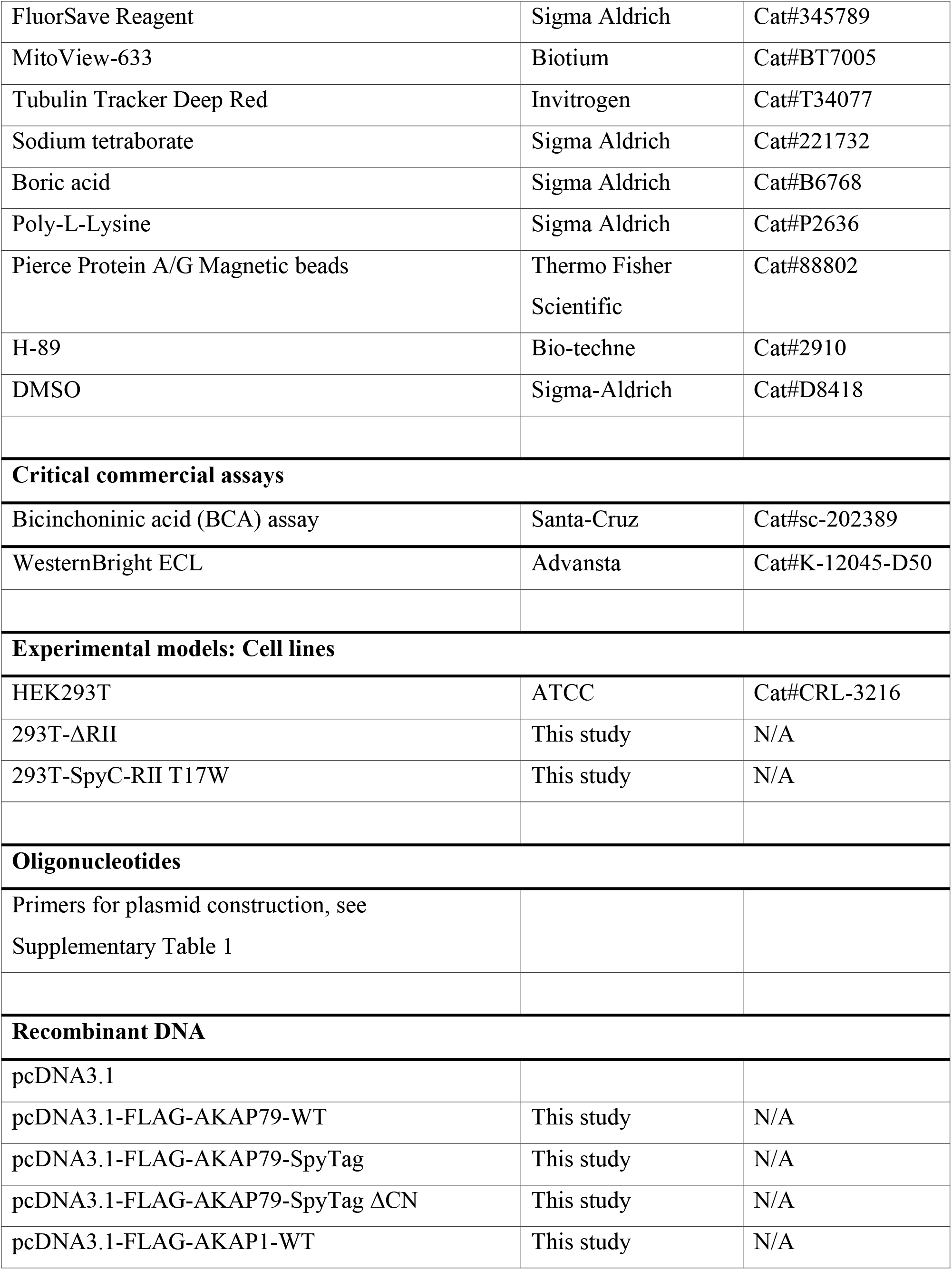
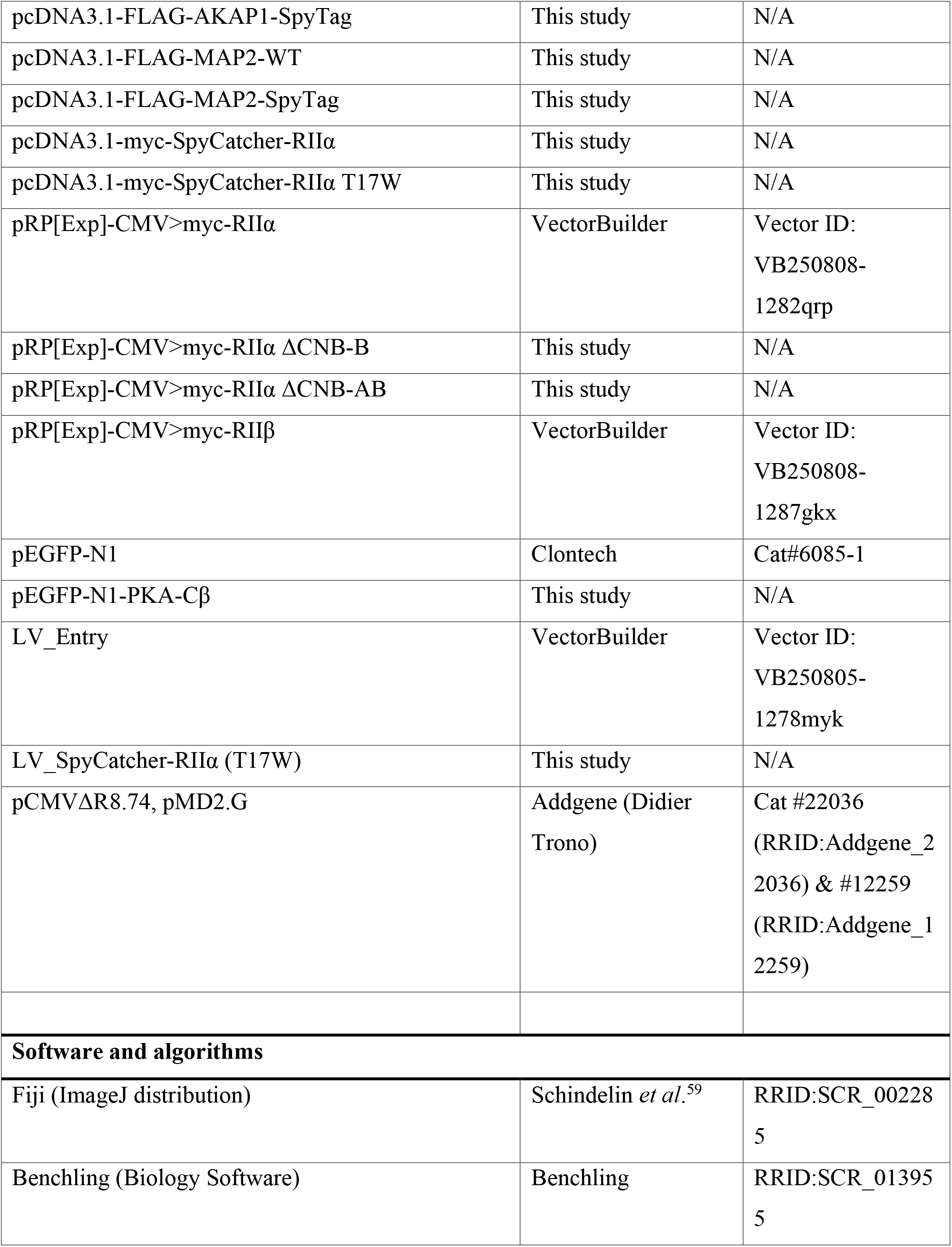
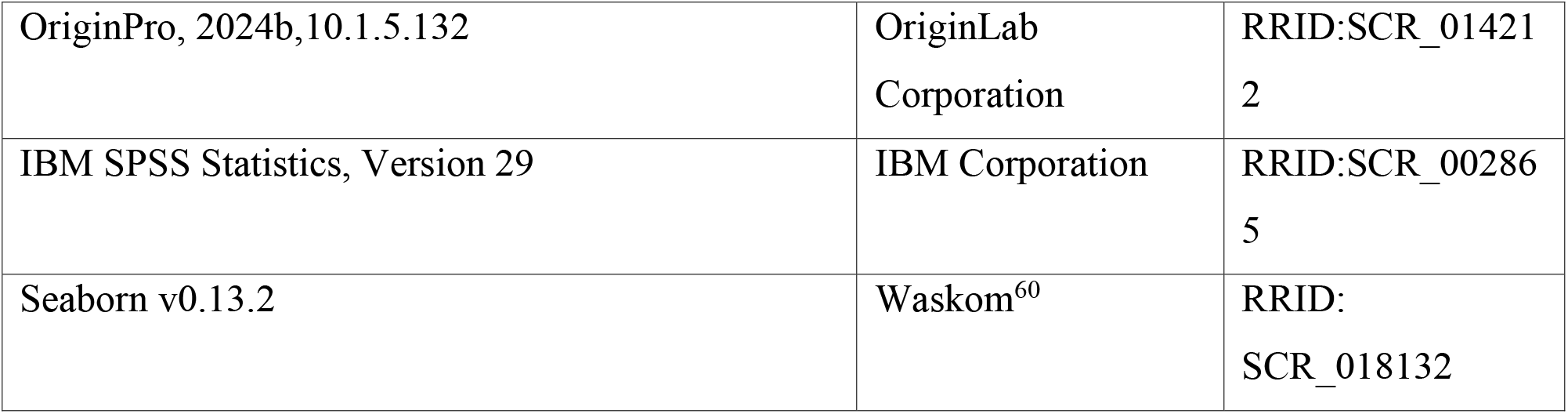

## Resource availability

### Lead contact

Further information and requests for resources and reagents should be directed to and will be fulfilled by the lead contact, MG.

### Materials availability

Plasmids generated in this study will be made available through Addgene and can be shared upon request.

### Data and code availability

- All data reported in this paper will be shared by the lead contact upon request.
- This paper does not report original code.
- Any additional information required to reanalyze the data reported in this paper is available from the lead contact upon request.

## Experimental Model & Study Participant Details

### Cell lines

HEK293T, ΔRII, and SpyC-RII cells were maintained in DMEM, high glucose, GlutaMAX Supplement, pyruvate (Gibco, Cat# 10569010), supplemented with 10% FBS and 1% penicillin/streptomycin. All cells were maintained at 37 °C with 5 % CO_2_. HEK293T cells were used for validating isopeptide-based PKA anchoring (**Fig. 1**). ΔRII cells were used for developing the SpyC-RIIα T17W cell line (**Fig. 2**), and to assess microtubule straightness (**Fig. 6**). The SpyC-RII cell line was used for all other experiments. HEK293T, ΔRII, and SpyC-RII cell lines were tested for mycoplasma contamination and were confirmed to be negative.

Primary hippocampal cultures were prepared from E18 Sprague-Dawley rats. Hippocampi were isolated and triturated with trypsin (0.025 %) before plating onto 13 mm coverslips, pre-treated with poly-L-lysine (0.1 mg/mL) at a density of 3×10^4^ per coverslip in DMEM containing 10 % heat-inactivated horse serum, and penicillin (40 U/mL)/streptomycin (50 µg/mL). Neurons were cultured in a 5 % CO_2_ humidified atmosphere at 37 °C. Two hours after seeding, the plating media was replaced with Neurobasal-A supplemented with 1 % B27, 0.5 % (v/v) GlutaMAX, 20 mM glucose, and penicillin (100 U/mL)/streptomycin (100 µg/mL). One-third of the media was replenished every 3-4 days after 7 days-in-vitro (DIV). Experiments were performed in accordance with the UK Animals Act 1986 and within University College London Animal Research guidelines overseen by the UCL Animal Welfare and Ethical Review Body under project code 14058.

## Method details

### DNA vector construction

FLAG-tagged AKAP constructs were assembled using the pcDNA3.1 vector backbone. pcDNA3.1-FLAG-AKAP79 WT was generated by Gibson assembly using primers 1FLAG_AKAP79_F/R and pcDNA3.1-2FLAG-AKAP79 template from a previous study^61^ whereas vectors containing AKAP1-FLAG WT (accession NM_003488.4) and MAP2-FLAG WT (accession NM_002374.3) were purchased from Genscript. AKAP variants containing the SpyTag sequence were generated using Gibson assembly. Positions 395-407 in AKAP79 were replaced with SpyTag using primers AKAP79_SpyTag_F/R, positions 363-375 in AKAP1 with primers AKAP1_SpyTag_F/R, and positions 86-98 in MAP2 with primers MAP2_SpyTag_F/R. ΔCN variants of AKAP79 were constructed by initially amplifying from template DNA containing the deletion Δ337-343^62^. For transient transfections, pcDNA3.1-myc-SpyCatcher-RIIα was assembled using primer pairs myc_SpyCatcher_Fwd/Rev for the SpyCatcher domain, and RIIα_Fwd/Rev for human RIIα. The resulting fusion protein contains a two-glycine linker between the minimal 114-amino acid SpyCatcher domain^25^ and RIIα. The T17W substitution was introduced using primers T17W_F/R. Mouse PKA Cβ (accession BC054533) was cloned N-terminal to EGFP in pEGFP-N1 (Clontech) using primers XhoI_Cbeta_M1 and Cbeta_F351_Glink_BamHI for expression of a fusion containing a glycine-rich linker (GGGGGSGGSGGSGGS) between Cβ and EGFP. Constructs for CMV promoter-driven transient expression of human myc-RIIα and myc-RIIβ were purchased from VectorBuilder. Truncated variants of RIIα lacking either the CNB-B domain or both CNB domains were generated by PCR with primer XbaI_RIIαTerm paired with either RIIα_DeltaB_XbaI or RIIα_DeltaAB_XbaI, digesting with XbaI prior to ligation and transformation. Arginine to lysine substitutions in RIIα were introduced by site-directed mutagenesis with primers RIIα_R217K_F/R and RIIα_R347K_F/R.

### Generation of ΔRII and SpyC-RII cell lines

To generate double RIIα/RIIβ knockout (ΔRII) cells, we employed CRISPR-Cas9 targeting. Guide sequences for PRKAR2A (5’GCCACATCCAGATCCCGCCG) and PRKAR2B (5’GATGAGCATCGAGATCCCGG) were designed in Benchling, and incorporated into full SafeEdit sgRNA sequences synthesised by Genscript (sgRNA-α and sgRNA-β). For ribonucleoprotein transfection, 2 separate 25 µL mixtures were prepared in Opti-MEM containing 0.5 μg of NLS-Cas9-NLS with 0.5 μg of either sgRNA-α or sgRNA-β. After 10 min incubation at 37 °C, the mixtures were combined then supplemented with 25 µL Opti-MEM containing 4 µL Lipofectamine-2000. The transfection mixture was incubated for a further 20 min and then added to HEK293T cells at ∼60% confluence in a 35 mm dish. Three days after transfection, cells were divided into 96-well plates at a density of 0.8 cells/well. Clones were expanded and screened for complete knockout of both RIIα and RIIβ expression using immunoblotting. SpyC-RII cells were generated using lentiviral infection of ΔRII cells. The lentiviral entry vector LV_Entry was ordered that includes the following elements between long-terminal repeats (5’ to 3’): CMV promoter; BamHI/XbaI entry sites; woodchuck hepatitis virus post-transcriptional regulatory element (WPRE); mPGK promoter driving blasticidin resistance gene. Myc-SpyCatcher-RIIα (T17W) sequence was amplified using primers BamHI_Myc_F and RIIα_term_XbaI and inserted into LV_Entry using BamHI and XbaI. The resulting LV_SpyCatcher-RIIα (T17W) vector was used to generate lentivirus by transfecting wild-type HEK293T cells with pCMVΔR8.74 packaging / pMD2.G envelope vectors. Virus was collected from the cell media 2 and 3 days after transfection, centrifuged for 5 min at 500 x *g*, filter sterilised, and snap frozen in liquid N_2_ for storage at -80 °C. For transduction, ΔRII cells at ∼ 60 % confluence were infected with a 1 in 50 dilution of virus in media. 24 hours later, the media was exchanged and supplemented with 5 µg/mL blasticidin to select for cells that had undergone viral integration. SpyC-RII clones were separated as before, and screened for SpyC-RII (T17W) expression at comparable levels to RIIα in wild-type HEK293T cells using immunoblotting.

### Transfections

Cells were generally transfected in 6-well plates, with 1.5 µg DNA / 6 µL Lipofectamine-2000 per well. For immunoprecipitation experiments, this was scaled up to 4 µg DNA / 16 µL Lipofectamine-2000 for transfection of cells in 10-cm plates. For live imaging experiments, DNA mixtures of 100 parts AKAPs to 1 part Cβ-EGFP were used to ensure that total C subunits did not exceed regulatory subunits. For EGFP and PKI-GFP co-transfections, DNA mixtures comprised 4 parts AKAP to 1 part EGFP/PKI-GFP. For co-transfections in ΔRII cells, MAP2 vectors were co-transfected with PKA-RII isoform expression vectors and pEGFP-N1 at a ratio of 89:10:1, respectively. For PLA, AKAP vectors were co-transfected with pEGFP-N1 in a 1:100 molar ratio, substituting the AKAP vector for pcDNA3.1 for the control condition. For protein extraction, cells were harvested 72 hours after transfection. For confocal imaging, cells were seeded onto coverslips treated with poly-L-lysine (0.25 mg/mL), or 35-mm glass bottom dishes, 2 days after transfection. Cells were then either fixed with PBS supplemented with 4 % paraformaldehyde (PFA)/3% sucrose, or imaged live, on the following day.

### Protein extraction & immunoprecipitation

The standard protein extraction procedure was to wash cells in PBS, then lyse in homogenisation buffer (30 mM Na Hepes pH 7.4, 0.5 mM EDTA, 150 mM NaCl, 1 mM Benzamidine, 1 % v/v Igepal-CA630, 0.25 % w/v sodium deoxycholate, 1 protease inhibitor cocktail/100 mL). For protein extraction prior to immunoblotting with anti-pRII antibody, cells were lysed in RIPA buffer (50 mM Tris pH 8, 150 mM NaCl, 0.5 % sodium deoxycholate, 0.1 % SDS, 1 % Igepal CA-630, 5 mM EDTA, 1 mM PMSF) supplemented with 1 PhosStop tablet per 10 mL, and 1 protease inhibitor cocktail tablet per 100 mL. In all cases, cells were sonicated at 20 kHz for 20 s and clarified by centrifugation at 20,000 x *g* for 20 min at 4 °C. Protein concentrations in extracts were determined by BCA assay. For immunoprecipitation, extracts were rotated overnight at 4 °C with 10 µL of Protein A/G magnetic beads (50 % slurry) and 2 µg of either rabbit anti-FLAG or rabbit IgG control antibody. On the following morning, beads were washed four times with homogenisation buffer before elution of bound proteins in 1x NuPAGE LDS sample buffer supplemented with 10 mM DTT for 5 min at 85 °C.

### Immunoblotting

Protein extracts were separated using NuPAGE 4-12 % Bis-Tris or 3-8 % Tris-acetate gels. Proteins were transferred onto nitrocellulose, blocked in 10 % milk/TBS-T for 1 hour before probing with primary antibodies in the same solution with the exception of anti-pRII antibody (5 % BSA/TBS-T). After overnight incubations, nitrocellulose membranes were washed (3 x 10 min incubations in 10 mL TBS-T), then incubated for 45 min with the relevant secondary antibody in 5% milk/TBS-T. Following three further washes as before, blots were developed with ECL substrate using an ImageQuant system. Densitometry was performed on immunoblot images using Fiji ^59^ within ImageJ (NIH). Antibody dilutions were as follows: anti-RIIα, 1 in 1000; anti-RIIβ, 1 in 1000; anti-C, 1 in 1000; anti-RI, 1 in 1000; anti-pRII, 1 in 2000; anti-FLAG-HRP, 1 in 1000; anti-FLAG, 1 in 1000; anti-myc, 1 in 2000; anti-GAPDH, 1 in 2500; anti-rabbit-HRP, 1 in 4000; anti-mouse-HRP, 1 in 4000.

### Staining and imaging of live cells

Nuclei were stained by adding Hoechst-33342 (1 μg/mL) to the cell media for 10 min. Cells were then washed and imaged in HEPES-buffered imaging solution (10 mM HEPES pH 7.4, 140 mM NaCl, 1 mM MgCl_2_, 5 mM KCl, 1 mM CaCl_2,_ 10 mM glucose). Cell membranes were labelled by 1 min incubation at 4 °C in cell maintenance media supplemented with 1 µg/mL WGA-CF633, before three washes in pre-chilled HEPES-buffered imaging solution. Cells stained with WGA-CF633 were imaged within 20 min of staining to minimize intracellular labelling. Mitochondria were stained with 20 nM MitoView-633 for 15 min at 37 °C in HEPES-buffered imaging solution before imaging. Microtubules were stained with Tubulin Tracker Deep Red (1 in 2000) in HEPES-buffered imaging solution for 30 min at 37 °C. In all cases, cells were washed three times with HEPES-buffered imaging solution before imaging. For WGA-CF633 staining, images were acquired within 20 min of staining to minimize intracellular accumulation of the stain. For PKA inhibition experiments, media was supplemented with either 10 µM H-89 or vehicle (DMSO) for 24 hours prior to and during imaging.

All imaging was performed using a Leica SP8 confocal microscope equipped with two high-sensitivity hybrid detectors, multiple lasers, and using a 63x (1.40 NA) HC Plan-Apo CS2 oil immersion objective to image cells in a chamber heated to 37 °C. C-GFP was imaged using 488 nm laser for excitation/500-550 nm emission filter. Far-red stains for the cell membrane, mitochondria, and microtubules were imaged using 633 nm laser/650-700 emission filter. Hoechst-33342 was imaged using 405 nm laser / 430-480 nm emission filter. WGA-CF633 signal was acquired using a scanning speed of 700 Hz and line average of 8; MitoView-633 with 1000 Hz scanning speed and line average of 8; Tubulin Tracker Deep Red at 400 Hz, line average of 4 and frame average of 2. The pinhole was set at 1 Airy unit for cell membrane and tubulin, with a z-interval of 0.5 μm whereas the pinhole was set at 0.45 Airy units and with z-stack intervals of 0.22 μm for mitochondria. All images were acquired at a resolution of 1024 x 1024.

### Proximity ligation assays

Cells were fixed three days after transfection in PBS supplemented with 4% PFA and 4 % sucrose for 15 mins at room temperature, washed three times with PBS, and permeabilized in PBS supplemented with 0.1 % Triton X-100 for 10 min. Following blocking in PBS supplemented with 10 % donkey serum for 60 min at room temperature, cells were incubated overnight at 4 °C in diluent buffer (PBS supplemented with 1 % donkey serum) containing the following primary antibodies: rabbit anti-FLAG (1:500); mouse anti-myc (1:500); chicken anti-GFP (1:500). On the following morning, cells were washed four times with diluent buffer before incubation with secondary PLA probes (anti-rabbit minus; anti-mouse plus) and anti-chicken Alexa Fluor 488 for 1 hour at 37 °C in a humidified chamber. Cells were then washed three times in Buffer A (10 mM Tris pH 7.4, 150 mM NaCl, 0.05 % Tween-20) before incubation in Duolink ligation solution for 30 min at 37 °C in a humidified chamber. After three further washes in Buffer A, amplification was performed with the Red Duolink reagent for 100 min at 37 °C in a humidified chamber. Finally, cells were washed twice in Buffer B (200 mM Tris pH 7.5, 100 mM NaCl), then once in 1 % Buffer B (diluted in water), before mounting on slides using ProLong Gold with DAPI, and sealing with nail varnish. Imaging was performed with the same microscope and objective used for live cell imaging, collecting Z-stacks in increments of 0.5 μm. DAPI was imaged using a 405 nm laser / 430-480 nm emission filter; GFP using 488/500-550 nm; and PLA puncta using 561/600-650 nm.

### Lentiviral transduction and imaging of primary hippocampal neurons

Lentiviruses for expression of shRIIα or scrambled RNA sequences were produced as described above using pFUGW-shRIIα and pFUGW-scrambled^14^ transfer vectors, respectively. Primary hippocampal neurons were transduced on DIV4 with a multiplicity of infection of 0.01 in conditioned neuronal media. The viral media was replaced 18 hours post-transduction with pre-conditioned neuronal media. Neurons were fixed on DIV14 using PBS supplemented with 4 % PFA and 4 % sucrose (15 min at room temperature). Neurons were permeabilized with 0.05 % Triton X-100 in PBS for 5 min then blocked for 1 hour in PBS supplemented with 10 % donkey serum. Neurons were incubated overnight at 4 °C with chicken anti-GFP antibody, diluted 1 in 1000 in PBS/1 % donkey serum. After washing, secondary incubation with anti-chicken Alexa Fluor 488 (diluted 1: 500 in PBS/1 % donkey serum) was performed for 1 hour in the dark. Coverslips were mounted onto glass slides using FluorSave mounting medium.

### Correlative Light and Electron Microscopy

SpyC-RII cells were co-transfected with C-GFP and either MAP2-WT or MAP2-SpyTag constructs. Forty-eight hours after transfection, cells were plated onto poly-L-lysine (0.1 mg/mL)-coated gridded glass-bottom dishes (Mattek, P35G-1.5-14-C-GRD). Live-cell imaging was performed 24 hours later at 37 °C using a Leica SP8 confocal microscope with cells in HEPES-based imaging buffer. Cells potentially suitable for EM were identified using C-GFP fluorescence (488 nm excitation laser, 500-550 nm emission filter), with grid coordinates mapped using brightfield microscopy and a 40x oil-immersion objective. Immediately following imaging, cells were fixed at 37 °C in PEM buffer (80 mM PIPES, pH 7.4; 2 mM MgCl₂; 1 mM EGTA) supplemented with 2 % EM-grade paraformaldehyde and 1.5 % EM-grade glutaraldehyde (TAAB Laboratories) for 30 min. Samples were then processed and embedded for electron microscopy as follows, with all stages occurring at room temperature unless otherwise stated: Samples were secondary fixed and stained in 1 % osmium tetroxide and 1.5 % potassium ferricyanide for 60 min in the dark, before washing in PEM buffer and subsequently further stained with 1 % tannic acid in PEM buffer for 45 min. Samples were then dehydrated by sequential washes in 70 % and 90 % ethanol in water, before two 100 % ethanol incubations for 10 min each. Samples were then infiltrated with epon resin via a 60 min incubation in 1:1 propylene oxide:epon mix, then two 60 min incubations in 100 % epon before inverting the coverslips onto pre polymerised epon stubs and baking overnight at 60 °C to polymerise the resin. Coverslips were removed via plunging in liquid nitrogen and target cells relocated using the grid relief pattern visible on the resin block surface. Ultrathin (70 nm) serial sections of each target cell were cut using an UC7 ultramicrotome (Leica) and a DiATOME 45° diamond knife, collected on formvar-coated 1 × 2 mm slot copper grids and stained with Reynolds lead citrate. Imaging was performed using a transmission electron microscope (Hitachi HT7800) at 100 kV and a Qedira CMOS camera (EMSIS).

### Image processing and analysis

All images were processed and analysed using Fiji Image (NIH). The JACoP plugin^63^ was used to generate Pearson’s correlation coefficients. Microtubule curvature was measured using the Kappa plugin, which uses squared distance minimisation to cubic B splines to quantify curving with output values corresponding to the inverse of the radius of curvature^41,64^. While analysis was performed on unfiltered images, representative images of microtubules are shown in figures after filtering using the tubeness plugin^65^ for clarity. Cell edge enrichment ratios were calculated by plotting C-GFP fluorescence intensity along line scans crossing the plasma membrane. The peak membrane-associated intensity was then divided by the intensity measured 1.5 µm inside the cell. Intensity values were averaged within 0.2 µm wedges along each line scan. Analysis of mitochondria was performed using the Mitochondria Analyzer Fiji plugin for image pre-processing, adaptive thresholding, and categorisation of mitochondria by size and shape^66^. For PLA, images were first masked according to the GFP channel (indicative of transfected cells), before calculating PLA signal area above a consistent threshold in each GFP positive cell^67^. For quantification of total cable length and branch point number in hippocampal neurons, individual neurons were manually reconstructed and traced using the Simple Neurite Tracer (SNT; v5.0.13) plugin^68^. For quantification of microtubule bundling characteristics in EM micrographs, bundles were defined as contiguous arrays of filaments running in parallel and separated by ≤140 nm. Inter-filament spacing was measured perpendicular to the filament axis and between filament centres. All measurements were performed on unprocessed micrographs.

### Experimental design, statistical testing and data visualisation

Each primary hippocampal culture was prepared from a pooled, mixed-sex population of approximately five E18 Sprague Dawley rat embryos from a single litter. For imaging analyses, individual cells were treated as the observational units and were pooled across at least three independent experiments. Each independent experiment comprised a separate culture preparation or, for cell-line experiments, a separate transfection and imaging session. The numbers of cells analysed are reported in the corresponding figures or figure legends. Where measurements were made on multiple structures within a cell, such as mitochondria or microtubules, these measurements were summarised at the cell level before statistical analysis unless otherwise stated.

Statistical analyses were performed using IBM SPSS Statistics v29, OriginPro 2024b, and Python. Data visualization was performed using OriginPro 2024 and the Seaborn Python library^60^. Data are presented as mean ± SEM, or as boxplots showing median and interquartile range, as indicated in figure legends. Immunoblot densitometry and PLA analyses in Figures 1 and 2 were analysed using unpaired two-tailed Student’s t-tests. Pearson’s correlation coefficient was used for image colocalization analyses. Where data were non-normally distributed as assessed by Shapiro-Wilk testing, ordinal, or strongly skewed, two-tailed Mann-Whitney U tests were used. Microtubule curvature and mitochondrial morphology datasets were analysed using two-tailed Mann-Whitney U tests, with Bonferroni correction applied for multiple pairwise comparisons where indicated. Neuronal cable length and branch point number were analysed using two-tailed Mann-Whitney U tests.

## Acknowledgements

We thank Didier Trono for plasmids pCMVΔR8.74 and pMD2.G (Addgene plasmids #22036 and #12259), which were essential for lentiviral production, and Hao Jiang for providing SpyCatcher coding sequences. We are grateful to Tao Peng and Minghao Lu for their support with pilot experiments. We thank colleagues in the Frances Brodsky and Patricia Salinas laboratories for helpful discussions on imaging, the staff of the University College London LMCB Electron Microscopy Core Facility (RRID:SCR_027340), and members of the EM of Cells tissues and Organisms Slack group for helpful advice and suggestions for microtubule preservation in EM preparations. This work was supported by a BBSRC project grant (BB/X008215/1) awarded to T.W.C. and M.G.G.

## Author contributions

Conceptualization, T.W.C., Y.L., and M.G.G.; Methodology, T.W.C., I.J.W., and M.G.G.; Investigation, all authors; Formal analysis, T.W.C. and M.G.G..; Writing – original draft, review and editing, T.W.C. and M.G.G. with input from all authors; Visualization, T.W.C. and M.G.G.; Supervision, M.G.G.; Funding acquisition, T.W.C. and M.G.G.

## Declaration of interests

The authors declare no competing interests.

## Notes

### Competing Interest Statement

The authors have declared no competing interest.

## References

1. Taylor, S.S., Wu, J., Bruystens, J.G.H., Del Rio, J.C., Lu, T.W., Kornev, A.P., and Ten Eyck, L.F. (2021). From structure to the dynamic regulation of a molecular switch: A journey over 3 decades. J Biol Chem 296, 100746. 10.1016/j.jbc.2021.100746.

2. Bers, D.M., Xiang, Y.K., and Zaccolo, M. (2019). Whole-Cell cAMP and PKA Activity are Epiphenomena, Nanodomain Signaling Matters. Physiology (Bethesda) 34, 240–249. 10.1152/physiol.00002.2019.

3. Kandel, E.R. (2012). The molecular biology of memory: cAMP, PKA, CRE, CREB-1, CREB-2, and CPEB. Mol Brain 5, 14. 10.1186/1756-6606-5-14.

4. Johnson, L.N. (2009). The regulation of protein phosphorylation. Biochem Soc Trans 37, 627–641. 10.1042/BST0370627.

5. Falcone, J.I., and Scott, J.D. (2025). The ascent of AKAPs, from architectural elements to kinase anchors: a perspective. The Biochemical journal 482, 485–498. 10.1042/BCJ20253085.

6. Gold, M.G., Lygren, B., Dokurno, P., Hoshi, N., McConnachie, G., Tasken, K., Carlson, C.R., Scott, J.D., and Barford, D. (2006). Molecular basis of AKAP specificity for PKA regulatory subunits. Mol Cell 24, 383–395.

7. Kinderman, F.S., Kim, C., von Daake, S., Ma, Y., Pham, B.Q., Spraggon, G., Xuong, N.H., Jennings, P.A., and Taylor, S.S. (2006). A dynamic mechanism for AKAP binding to RII isoforms of cAMP-dependent protein kinase. Mol Cell 24, 397–408.

8. Dell’Acqua, M.L., Faux, M.C., Thorburn, J., Thorburn, A., and Scott, J.D. (1998). Membrane-targeting sequences on AKAP79 bind phosphatidylinositol-4, 5-bisphosphate. EMBO J 17, 2246–2260. 10.1093/emboj/17.8.2246.

9. Lewis, S.A., Wang, D.H., and Cowan, N.J. (1988). Microtubule-associated protein MAP2 shares a microtubule binding motif with tau protein. Science 242, 936–939. 10.1126/science.3142041.

10. Huang, L.J., Wang, L., Ma, Y., Durick, K., Perkins, G., Deerinck, T.J., Ellisman, M.H., and Taylor, S.S. (1999). NH2-Terminal targeting motifs direct dual specificity A-kinase-anchoring protein 1 (D-AKAP1) to either mitochondria or endoplasmic reticulum. The Journal of cell biology 145, 951–959. 10.1083/jcb.145.5.951.

11. Gold, M.G. (2019). Swimming regulations for protein kinase A catalytic subunit. Biochem Soc Trans 47, 1355–1366. 10.1042/BST20190230.

12. Zhang, P., Knape, M.J., Ahuja, L.G., Keshwani, M.M., King, C.C., Sastri, M., Herberg, F.W., and Taylor, S.S. (2015). Single Turnover Autophosphorylation Cycle of the PKA RIIbeta Holoenzyme. PLoS Biol 13, e1002192. 10.1371/journal.pbio.1002192.

13. Walker-Gray, R., Stengel, F., and Gold, M.G. (2017). Mechanisms for restraining cAMP-dependent protein kinase revealed by subunit quantitation and cross-linking approaches. Proc Natl Acad Sci U S A 114, 10414–10419. 10.1073/pnas.1701782114.

14. Church, T.W., Tewatia, P., Hannan, S., Antunes, J., Eriksson, O., Smart, T.G., Hellgren Kotaleski, J., and Gold, M.G. (2021). AKAP79 enables calcineurin to directly suppress protein kinase A activity. Elife 10. 10.7554/eLife.68164.

15. Aye, T.T., Scholten, A., Taouatas, N., Varro, A., Van Veen, T.A., Vos, M.A., and Heck, A.J. (2010). Proteome-wide protein concentrations in the human heart. Molecular bioSystems 6, 1917–1927. 10.1039/c004495d.

16. Tullis, J.E., Larsen, M.E., Rumian, N.L., Freund, R.K., Boxer, E.E., Brown, C.N., Coultrap, S.J., Schulman, H., Aoto, J., Dell’Acqua, M.L., and Bayer, K.U. (2023). LTP induction by structural rather than enzymatic functions of CaMKII. Nature 621, 146–153. 10.1038/s41586-023-06465-y.

17. Araki, Y., Rajkovich, K.E., Gerber, E.E., Gamache, T.R., Johnson, R.C., Tran, T.H.N., Liu, B., Zhu, Q., Hong, I., Kirkwood, A., and Huganir, R. (2024). SynGAP regulates synaptic plasticity and cognition independently of its catalytic activity. Science 383, eadk1291. 10.1126/science.adk1291.

18. Carr, D.W., Hausken, Z.E., Fraser, I.D., Stofko-Hahn, R.E., and Scott, J.D. (1992). Association of the type II cAMP-dependent protein kinase with a human thyroid RII-anchoring protein. Cloning and characterization of the RII-binding domain. J Biol Chem 267, 13376–13382.

19. Carlson, C.R., Lygren, B., Berge, T., Hoshi, N., Wong, W., Tasken, K., and Scott, J.D. (2006). Delineation of type I protein kinase A-selective signaling events using an RI anchoring disruptor. J Biol Chem 281, 21535–21545.

20. Wang, Y., Ho, T.G., Bertinetti, D., Neddermann, M., Franz, E., Mo, G.C., Schendowich, L.P., Sukhu, A., Spelts, R.C., Zhang, J., et al. (2014). Isoform-selective disruption of AKAP-localized PKA using hydrocarbon stapled peptides. ACS chemical biology 9, 635–642. 10.1021/cb400900r.

21. Gold, M.G., Fowler, D.M., Means, C.K., Pawson, C.T., Stephany, J.J., Langeberg, L.K., Fields, S., and Scott, J.D. (2013). Engineering A-kinase anchoring protein (AKAP)-selective regulatory subunits of protein kinase A (PKA) through structure-based phage selection. J Biol Chem 288, 17111–17121. 10.1074/jbc.M112.447326.

22. Hagan, R.M., Bjornsson, R., McMahon, S.A., Schomburg, B., Braithwaite, V., Buhl, M., Naismith, J.H., and Schwarz-Linek, U. (2010). NMR spectroscopic and theoretical analysis of a spontaneously formed Lys-Asp isopeptide bond. Angew Chem Int Ed Engl 49, 8421–8425. 10.1002/anie.201004340.

23. Zakeri, B., Fierer, J.O., Celik, E., Chittock, E.C., Schwarz-Linek, U., Moy, V.T., and Howarth, M. (2012). Peptide tag forming a rapid covalent bond to a protein, through engineering a bacterial adhesin. Proc Natl Acad Sci U S A 109, E690–697. 10.1073/pnas.1115485109.

24. Hatlem, D., Trunk, T., Linke, D., and Leo, J.C. (2019). Catching a SPY: Using the SpyCatcher-SpyTag and Related Systems for Labeling and Localizing Bacterial Proteins. Int J Mol Sci 20. 10.3390/ijms20092129.

25. Li, L., Fierer, J.O., Rapoport, T.A., and Howarth, M. (2014). Structural analysis and optimization of the covalent association between SpyCatcher and a peptide Tag. J Mol Biol 426, 309–317. 10.1016/j.jmb.2013.10.021.

26. Herberg, F.W., Maleszka, A., Eide, T., Vossebein, L., and Tasken, K. (2000). Analysis of A-kinase anchoring protein (AKAP) interaction with protein kinase A (PKA) regulatory subunits: PKA isoform specificity in AKAP binding. J Mol Biol 298, 329–339. 10.1006/jmbi.2000.3662.

27. Hardy, J.C., Pool, E.H., Bruystens, J.G.H., Zhou, X., Li, Q., Zhou, D.R., Palay, M., Tan, G., Chen, L., Choi, J.L.C., et al. (2024). Molecular determinants and signaling effects of PKA RIalpha phase separation. Mol Cell 84, 1570–1584 e1577. 10.1016/j.molcel.2024.03.002.

28. Thomas, R., Jacoby, P.S., De Faveri, C., Derieux, C., Liebing, A.D., Melkes, B., Martini, H.J., Bermudez, M., Staubert, C., Lohse, M.J., et al. (2026). Ligand-specific activation trajectories dictate GPCR signalling in cells. Nature 650, 1053–1062. 10.1038/s41586-025-09963-3.

29. Willoughby, D., Wong, W., Schaack, J., Scott, J.D., and Cooper, D.M. (2006). An anchored PKA and PDE4 complex regulates subplasmalemmal cAMP dynamics. EMBO J 25, 2051–2061. 10.1038/sj.emboj.7601113.

30. Alam, M.S. (2022). Proximity Ligation Assay (PLA). Methods Mol Biol 2422, 191–201. 10.1007/978-1-0716-1948-3_13.

31. Tillo, S.E., Xiong, W.H., Takahashi, M., Miao, S., Andrade, A.L., Fortin, D.A., Yang, G., Qin, M., Smoody, B.F., Stork, P.J.S., and Zhong, H. (2017). Liberated PKA Catalytic Subunits Associate with the Membrane via Myristoylation to Preferentially Phosphorylate Membrane Substrates. Cell reports 19, 617–629. 10.1016/j.celrep.2017.03.070.

32. Xiong, W., Qin, M., and Zhong, H. (2024). PKA regulation of neuronal function requires the dissociation of catalytic subunits from regulatory subunits. Elife 13. 10.7554/eLife.93766.

33. Dittmer, P.J., Dell’Acqua, M.L., and Sather, W.A. (2014). Ca2+/calcineurin-dependent inactivation of neuronal L-type Ca2+ channels requires priming by AKAP-anchored protein kinase A. Cell reports 7, 1410–1416. 10.1016/j.celrep.2014.04.039.

34. Cochrane, V.A., Yang, Z., Dell’Acqua, M.L., and Shyng, S.L. (2021). AKAP79/150 coordinates leptin-induced PKA signaling to regulate K(ATP) channel trafficking in pancreatic beta-cells. J Biol Chem 296, 100442. 10.1016/j.jbc.2021.100442.

35. Theurkauf, W.E., and Vallee, R.B. (1982). Molecular characterization of the cAMP-dependent protein kinase bound to microtubule-associated protein 2. J Biol Chem 257, 3284–3290.

36. Zhong, H., Sia, G.M., Sato, T.R., Gray, N.W., Mao, T., Khuchua, Z., Huganir, R.L., and Svoboda, K. (2009). Subcellular dynamics of type II PKA in neurons. Neuron 62, 363–374. 10.1016/j.neuron.2009.03.013.

37. Harada, A., Teng, J., Takei, Y., Oguchi, K., and Hirokawa, N. (2002). MAP2 is required for dendrite elongation, PKA anchoring in dendrites, and proper PKA signal transduction. The Journal of cell biology 158, 541–549. 10.1083/jcb.200110134.

38. Leterrier, J.F., Kurachi, M., Tashiro, T., and Janmey, P.A. (2009). MAP2-mediated in vitro interactions of brain microtubules and their modulation by cAMP. Eur Biophys J 38, 381–393. 10.1007/s00249-008-0381-1.

39. Itoh, T.J., Hisanaga, S., Hosoi, T., Kishimoto, T., and Hotani, H. (1997). Phosphorylation states of microtubule-associated protein 2 (MAP2) determine the regulatory role of MAP2 in microtubule dynamics. Biochemistry 36, 12574–12582. 10.1021/bi962606z.

40. Lyu, J., DeMarco, A.G., Sweet, R.A., and Grubisha, M.J. (2025). MAP2 phosphorylation: mechanisms, functional consequences, and emerging insights. Front Cell Neurosci 19, 1610371. 10.3389/fncel.2025.1610371.

41. Mary, H., and Brouhard, G.J. (2019). Kappa: Analysis of Curvature in Biological Image Data using B-splines. bioRxiv. 10.1101/852772.

42. Huang, L.J., Durick, K., Weiner, J.A., Chun, J., and Taylor, S.S. (1997). Identification of a novel protein kinase A anchoring protein that binds both type I and type II regulatory subunits. J Biol Chem 272, 8057–8064. 10.1074/jbc.272.12.8057.

43. Gabrovsek, L., Collins, K.B., Aggarwal, S., Saunders, L.M., Lau, H.T., Suh, D., Sancak, Y., Trapnell, C., Ong, S.E., Smith, F.D., and Scott, J.D. (2020). A-kinase-anchoring protein 1 (dAKAP1)-based signaling complexes coordinate local protein synthesis at the mitochondrial surface. J Biol Chem 295, 10749–10765. 10.1074/jbc.RA120.013454.

44. Yu, R., Liu, T., Ning, C., Tan, F., Jin, S.B., Lendahl, U., Zhao, J., and Nister, M. (2019). The phosphorylation status of Ser-637 in dynamin-related protein 1 (Drp1) does not determine Drp1 recruitment to mitochondria. J Biol Chem 294, 17262–17277. 10.1074/jbc.RA119.008202.

45. Chang, C.R., and Blackstone, C. (2007). Cyclic AMP-dependent protein kinase phosphorylation of Drp1 regulates its GTPase activity and mitochondrial morphology. J Biol Chem 282, 21583–21587. 10.1074/jbc.C700083200.

46. Bridges, D., MacDonald, J.A., Wadzinski, B., and Moorhead, G.B. (2006). Identification and characterization of D-AKAP1 as a major adipocyte PKA and PP1 binding protein. Biochemical and biophysical research communications 346, 351–357. 10.1016/j.bbrc.2006.05.138.

47. Burdyga, A., Surdo, N.C., Monterisi, S., Di Benedetto, G., Grisan, F., Penna, E., Pellegrini, L., Zaccolo, M., Bortolozzi, M., Swietach, P., et al. (2018). Phosphatases control PKA-dependent functional microdomains at the outer mitochondrial membrane. Proc Natl Acad Sci U S A 115, E6497–E6506. 10.1073/pnas.1806318115.

48. Conca, F., Bayburtlu, D.K., Vismara, M., Surdo, N.C., Tavoni, A., Nogara, L., Sarra, A., Ciciliot, S., Di Benedetto, G., Iannucci, L.F., and Lefkimmiatis, K. (2025). Phosphatases Control the Duration and Range of cAMP/PKA Microdomains. Function (Oxf) 6. 10.1093/function/zqaf007.

49. Mehta, S., Zhang, Y., Roth, R.H., Zhang, J.F., Mo, A., Tenner, B., Huganir, R.L., and Zhang, J. (2018). Single-fluorophore biosensors for sensitive and multiplexed detection of signalling activities. Nature cell biology 20, 1215–1225. 10.1038/s41556-018-0200-6.

50. McNicholl, E.T., Das, R., SilDas, S., Taylor, S.S., and Melacini, G. (2010). Communication between tandem cAMP binding domains in the regulatory subunit of protein kinase A-Ialpha as revealed by domain-silencing mutations. J Biol Chem 285, 15523–15537. 10.1074/jbc.M110.105783.

51. Brown, S.H., Cheng, C.Y., Saldanha, S.A., Wu, J., Cottam, H.B., Sankaran, B., and Taylor, S.S. (2013). Implementing fluorescence anisotropy screening and crystallographic analysis to define PKA isoform-selective activation by cAMP analogs. ACS chemical biology 8, 2164–2172. 10.1021/cb400247t.

52. Bastidas, A.C., Deal, M.S., Steichen, J.M., Keshwani, M.M., Guo, Y., and Taylor, S.S. (2012). Role of N-terminal myristylation in the structure and regulation of cAMP-dependent protein kinase. J Mol Biol 422, 215–229. 10.1016/j.jmb.2012.05.021.

53. Xiong, W.H., Qin, M., and Zhong, H. (2021). Myristoylation alone is sufficient for PKA catalytic subunits to associate with the plasma membrane to regulate neuronal functions. Proc Natl Acad Sci U S A 118. 10.1073/pnas.2021658118.

54. Sanchez, C., Diaz-Nido, J., and Avila, J. (2000). Phosphorylation of microtubule-associated protein 2 (MAP2) and its relevance for the regulation of the neuronal cytoskeleton function. Prog Neurobiol 61, 133–168. 10.1016/s0301-0082(99)00046-5.

55. Khuchua, Z., Wozniak, D.F., Bardgett, M.E., Yue, Z., McDonald, M., Boero, J., Hartman, R.E., Sims, H., and Strauss, A.W. (2003). Deletion of the N-terminus of murine map2 by gene targeting disrupts hippocampal ca1 neuron architecture and alters contextual memory. Neuroscience 119, 101–111. 10.1016/s0306-4522(03)00094-0.

56. Fujioka, T., Fujioka, A., and Duman, R.S. (2004). Activation of cAMP signaling facilitates the morphological maturation of newborn neurons in adult hippocampus. J Neurosci 24, 319–328. 10.1523/JNEUROSCI.1065.03.2004.

57. Gorshkov, K., Mehta, S., Ramamurthy, S., Ronnett, G.V., Zhou, F.Q., and Zhang, J. (2017). AKAP-mediated feedback control of cAMP gradients in developing hippocampal neurons. Nature chemical biology 13, 425–431. 10.1038/nchembio.2298.

58. Kim, Y., Jang, Y.N., Kim, J.Y., Kim, N., Noh, S., Kim, H., Queenan, B.N., Bellmore, R., Mun, J.Y., Park, H., et al. (2020). Microtubule-associated protein 2 mediates induction of long-term potentiation in hippocampal neurons. FASEB J 34, 6965–6983. 10.1096/fj.201902122RR.

59. Schindelin, J., Arganda-Carreras, I., Frise, E., Kaynig, V., Longair, M., Pietzsch, T., Preibisch, S., Rueden, C., Saalfeld, S., Schmid, B., et al. (2012). Fiji: an open-source platform for biological-image analysis. Nat Methods 9, 676–682. 10.1038/nmeth.2019.

60. Waskom, M.L. (2021). seaborn: statistical data visualization. Journal of Open Source Software 6, 3021. 10.21105/joss.03021.

61. Patel, N., Stengel, F., Aebersold, R., and Gold, M.G. (2017). Molecular basis of AKAP79 regulation by calmodulin. Nature communications 8, 1681. 10.1038/s41467-017-01715-w.

62. Gold, M.G., Stengel, F., Nygren, P.J., Weisbrod, C.R., Bruce, J.E., Robinson, C.V., Barford, D., and Scott, J.D. (2011). Architecture and dynamics of an A-kinase anchoring protein 79 (AKAP79) signaling complex. Proc Natl Acad Sci U S A 108, 6426–6431.

63. Bolte, S., and Cordelieres, F.P. (2006). A guided tour into subcellular colocalization analysis in light microscopy. J Microsc 224, 213–232. 10.1111/j.1365-2818.2006.01706.x.

64. Sasaki, T., Saito, K., Inoue, D., Serk, H., Sugiyama, Y., Pesquet, E., Shimamoto, Y., and Oda, Y. (2023). Confined-microtubule assembly shapes three-dimensional cell wall structures in xylem vessels. Nature communications 14, 6987. 10.1038/s41467-023-42487-w.

65. Sato, Y., Nakajima, S., Shiraga, N., Atsumi, H., Yoshida, S., Koller, T., Gerig, G., and Kikinis, R. (1998). Three-dimensional multi-scale line filter for segmentation and visualization of curvilinear structures in medical images. Med Image Anal 2, 143–168. 10.1016/s1361-8415(98)80009-1.

66. Chaudhry, A., Shi, R., and Luciani, D.S. (2020). A pipeline for multidimensional confocal analysis of mitochondrial morphology, function, and dynamics in pancreatic beta-cells. Am J Physiol Endocrinol Metab 318, E87–E101. 10.1152/ajpendo.00457.2019.

67. Hegazy, M., Cohen-Barak, E., Koetsier, J.L., Najor, N.A., Arvanitis, C., Sprecher, E., Green, K.J., and Godsel, L.M. (2020). Proximity Ligation Assay for Detecting Protein-Protein Interactions and Protein Modifications in Cells and Tissues in Situ. Curr Protoc Cell Biol 89, e115. 10.1002/cpcb.115.

68. Arshadi, C., Gunther, U., Eddison, M., Harrington, K.I.S., and Ferreira, T.A. (2021). SNT: a unifying toolbox for quantification of neuronal anatomy. Nat Methods 18, 374–377. 10.1038/s41592-021-01105-7.

