## Supplemental Information for "PKA can adopt structural or enzymatic roles depending on its anchoring site"

**Document S1**. Figures S1, S2, S3 and Table S1.

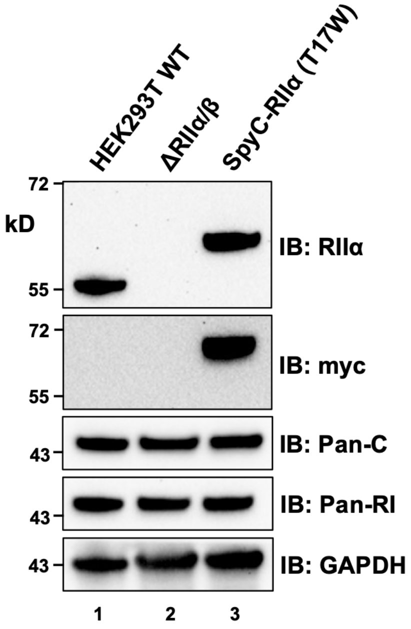


**Figure S1. PKA subunit expression levels in HEK293T cell line variants**.
Expression levels of RIIα, myc-SpyCatcher-RIIα T17W, C subunit, and RI subunit levels were compared in HEK293T WT cell extracts (lane 1), along with extracts from ΔRII (lane 2) and SpyC-RII (lane 3) HEK293T variant extracts. Anti-GAPDH immunoblotting was included as a loading control. The immunoblots did not reveal any compensatory changes in RI or C subunit expression levels in the HEK293T variant cell lines.

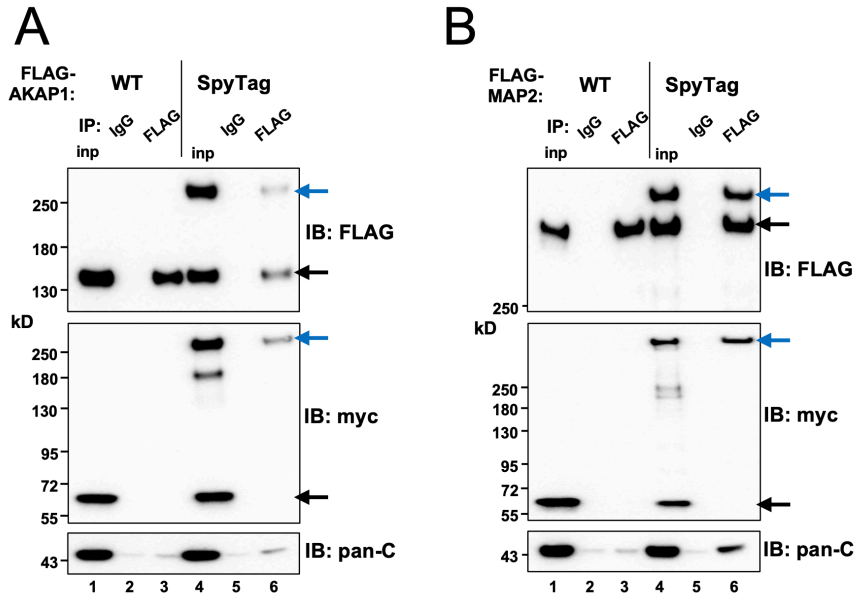


**Figure S2. Immunoprecipitation experiments to validate isopeptide anchoring to AKAP1 and MAP2**.

Co-immunoprecipitation experiments are shown for protein extracts from SpyC-RII cells transfected with WT and SpyTag variants of either (**A**) FLAG-AKAP1 or (**B**) FLAG-MAP2. Proteins were immunoprecipitated with either control IgG (lanes 2 & 5) or anti-FLAG (lanes 3 & 6) antibody. IP of FLAG-tagged AKAPs is shown in the top sub-panels; co-IP of myc-SpyCatcher-RII subunits in the middle sub-panels; and co-IP of C subunits in the lower sub-panels. Unconjugated and conjugated proteins are indicated by black and blue arrows, respectively.

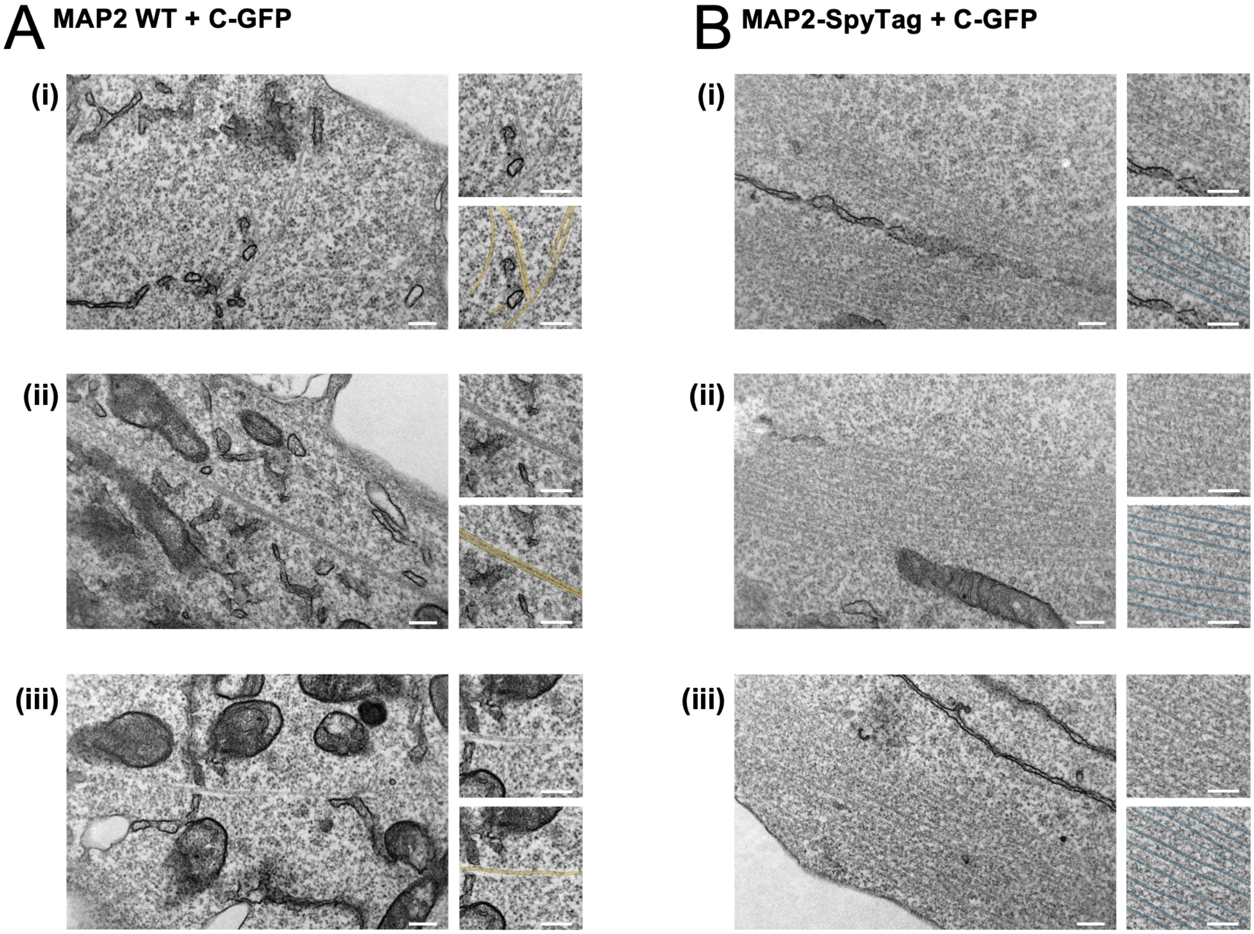


**Figure S3. Additional electron micrographs supporting the microtubule-bundling analysis in Fig. 7A and Fig. 7D.**
(A) Electron micrographs from SpyC-RII cells co-expressing C-GFP and MAP2-WT, corresponding to the boxed regions i–iii shown in Fig. 7A. (B) Electron micrographs from SpyC-RII cells co-expressing C-GFP and MAP2-SpyTag, corresponding to the boxed regions i–iii shown in Fig. 7D. Each row shows a representative electron micrograph together with enlarged unannotated and annotated views of the same region. Microtubule filaments are highlighted in orange for MAP2-WT and cyan for MAP2-SpyTag. Scale bars: 200 nm.

**Table S1. Oligonucleotide sequences used for vector construction, related to STAR methods.**

| Primer Name | Sequence (5' to 3') |
|---|---|
| Myc_SpyCatcher_Fwd | TTGGTACCATGGAGCAGAAACTCATCTCTGAAGAGGATCTGGCAACCCATATTAAATTTAGCAAACG |
| SpyCatcher_Rev | TGGTCGACACCTTTGGTTGCTTTACCATTAACG |
| RIIα_Fwd | GGCGGATCCCACATCCAGATCCCGCCG |
| RIIα_Rev | TTCCTCGAGCTACTGCCCGAGGTTGCC |
| BamHI_Myc_F | AAATGGATCCGCCACCATGGAGCAGAAACTCATC |
| RIIα_term_XbaI | AAAATCTAGACTACTGCCCGAGGTTG |
| T17W_F | TGGGTGGAGGTGCTGCGACAGC |
| T17W_R | GTAGCCCTGCAGCAGCTCCG |
| 1FLAG_AKAP79_F | CTACAAAGATGATGACGATAAAATGGAAACCACAATTTCAG |
| 1FLAG_AKAP79_R | CGTCATCATCTTTGTAGTCCATGGTGGGATCCGAGC |
| SV40_Gibson_F | GTATGCAAAGCATGCATCTCAATTAGTCAG |
| SV40_Gibson_R | CTGACTAATTGAGATGCATGCTTTGCATAC |
| AKAP79_SpyTag_F | GAAGCACATATTGTTATGGTTGATGCGTATAAACCGACCAAGATAGAACAGCTGGTTAATGAAATGGCC |
| AKAP79_SpyTag_R | TATCTTGGTCGGTTTATACGCATCAACCATAACAATATGTGCTTCAATTAAGAGTGTTTCATATTGTTC |
| MAP2_SpyTag_F | GAAGCACATATTGTTATGGTTGATGCGTATAAACCGACCAAGACTGCTGAGGCTGTAGCAGTCCTGAAAGG |
| MAP2_SpyTag_R | AGTCTTGGTCGGTTTATACGCATCAACCATAACAATATGTGCTTCTCTGTCAGCTGAGGTCAGCTCTCC |
| AKAP1_SpyTag_F | GCTCATATTGTTATGGTTGATGCGTATAAACCGACCAAGACCGAACAGGTGCTGGCCACCACGGTTGGC |
| AKAP1_SpyTag_R | CTTGGTCGGTTTATACGCATCAACCATAACAATATGAGCCCGCTTAATCTCCTCATTTCTATC |
| XhoI_Cbeta_M1wt | ATCGCTCTCGAGCACCATGGGGAACACTGCGATCGCCAAG |
| Cbeta_F351_Glink_BamHI | ATAATGGATCCGAGCCGCTGCCACCAGAGCCACCGCTACCGCCGCCACCGCCAAATTCACAAAATTCCTTTCCACATTTTTC |

| | |
|---|---|
| RIIα_DeltaAB_XbaI | AAATATCTAGACTAGAGAATATCTTTGCAAGCTTCCTG |
| RIIα_DeltaB_XbaI | AAATATCTAGACTAGGGCACAGACTCAATAAATGATTCAAAC |
| XbaI_RIIαTerm | GTCGACTCTAGAACCCAGCTTTC |
| RIIα_R217K_F | CAACACCCCGAAAGCTGCTACCATTGTTG |
| RIIα_R217K_R | GCAGCTTTCGGGGTGTTGTACATCAG |
| RIIα_R347K_F | CAACAAACCCAAAGCTGCCTCAGCTTATGC |
| RIIα_R347K_R | CTGAGGCAGCTTTGGGTTTGTTGGTGACCAG |
